# Structural basis of anticancer drug recognition and amino acid transport by LAT1

**DOI:** 10.1101/2023.12.03.567112

**Authors:** Yongchan Lee, Chunhuan Jin, Ryuichi Ohgaki, Minhui Xu, Satoshi Ogasawara, Rangana Warshamanage, Keitaro Yamashita, Garib Murshudov, Osamu Nureki, Takeshi Murata, Yoshikatsu Kanai

## Abstract

LAT1 (SLC7A5) transports large neutral amino acids and their derivatives across the plasma membrane and plays pivotal roles in cancer cell proliferation, immune response and drug delivery across the blood-brain barrier. Despite recent advances in structural understanding of LAT1, how it discriminates substrates and inhibitors including the clinically relevant anticancer drugs remains elusive. Here we report six structures of LAT1, captured in three different conformations and bound with diverse bioactive ligands, elucidating its substrate transport and inhibitory mechanisms. JPH203, also known as nanvuranlat or KYT-0353 and currently in clinical trials as an anticancer drug, binds to the wide-open substrate-binding pocket of LAT1. It adopts a U-shaped conformer, with its amino-phenylbenzoxazol moiety pushing against transmembrane helix 3 (TM3), bending TM10 and arresting the transporter in the outward-facing conformation. In contrast, the physiological substrate L-Phe does not exhibit such inhibitory interactions, whereas melphalan, a slow substrate, poses steric hindrance in the pocket, explaining its inhibitory activity. Unexpectedly, the “classical” system L inhibitor BCH induces an occluded state, a key structural intermediate required for substrate transport. *Trans* stimulation assays show that BCH facilitates transporter turnover and is therefore a transportable substrate. These findings provide a structural framework for the intricate mechanisms of substrate recognition and inhibition of LAT1, paving the way for developing more specific and effective drugs against it.

## Introduction

System L is the major amino acid transport system that supplies large neutral amino acids into cells (*1*, *2*) (Fig. 1a). Among four molecular species constituting system L (LAT1–4), LAT1 is unique in its wide substrate specificity (*3–5*), which ranges from neutral amino acids such as L-Leu and L-Phe to much bulkier derivatives such as thyroid hormones (e.g. T_4_ and T_3_) and an alkylating agent melphalan (Fig. 1b). Since LAT1 is upregulated in various types of cancer, its inhibitors are known as potent anti-tumor agents (*6*) (Fig. 1a), its radiotracers are used in the positron emission tomography (PET) and the single photon emission computed tomography (SPECT) for cancer imaging (*7*, *8*), and the boronated substrates are employed for boron neutron capture therapy (BNCT) for cancer treatment (*7*, *8*). LAT1 is also responsible for the delivery of amino acid drugs and pro-drugs across the blood-brain barrier (BBB), such as gabapentin and L-DOPA (Fig. 1b), prescribed for epilepsy (*9*) and Parkinson’s disease (*10*), respectively.

**Figure 1.**
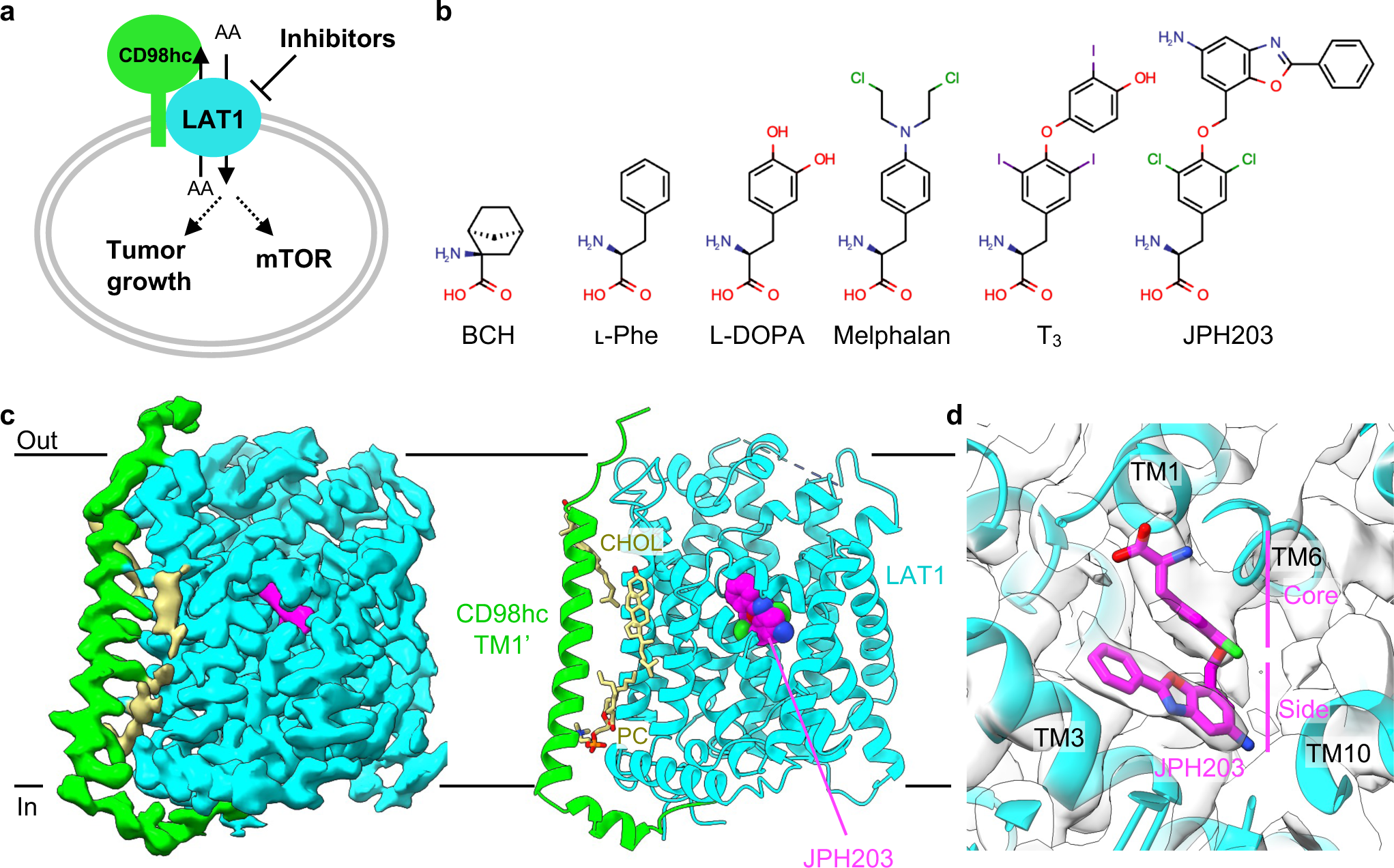
Structure of LAT1 bound to JPH203. **a)** Schematic representation of the LAT1 functions in cancer cells. **b)** Chemical structures of selected LAT1 substrates and inhibitors. **c)** Cryo-EM map and the overall structure of LAT1 bound to JPH203. The map of the transmembrane domain after local refinement is shown. Nanodisc densities are not displayed for clarity. **d)** Close-up view of the JPH203-binding site overlayed with the map.

JPH203 (Fig. 1b) is a high-affinity inhibitor of LAT1 with a sub-micromolar IC_50_ and no detectable inhibition on LAT2 (*11*). With such outstanding selectivity and potency, JPH203 has been proved effective against different types of cancers (*12*, *13*) and successfully completed Phase I and II clinical trials as a first-in-class drug against biliary tract cancer (*14*, *15*). Although its structural design is inspired by T_3_, JPH203 has a bulkier hydrophobic side chain, which may be responsible for its high selectivity and affinity (Fig. 1b). 2-aminobicyclo-(2,2,1)-heptane-2-carboxylic acid, also known as BCH, is a “classical” system L inhibitor with broad specificity towards both LAT1 and LAT2, as well as other system L (*1*). BCH has a bicyclic norbornane moiety and is significantly smaller than other Tyr-based inhibitors (Fig. 1b).

A recent structural study of LAT1 showed that JPH203 did not bind to their purified LAT1 (*16*), presumably due to the presence of detergent, which hindered the structural understanding of JPH203. Additionally, the same study reported a LAT1 structure with a BCH ligand modelled; however, the ligand density was very weak and indistinguishable from the apo map, and caution is required when interpreting this model (Fig. S1, difference maps). Therefore, the inhibition mechanism by BCH remains to be elucidated. More recently, JX molecules, which are bicyclic meta-Tyr derivatives, were reported as high affinity inhibitors and the associated cryo-EM structures revealed how these compounds bind to the outward-occluded conformations of LAT1 (*17*). This was the first structural demonstration of system L inhibition, but the different core structures of JX inhibitors compared to JPH203 and other amino acid-like compounds have hampered the understanding of how the well-known system L inhibitors bind to and inhibit LAT1.

Here, we employ the lipid nanodisc system and electron cryo-microscopy (cryo-EM) to study the structure of LAT1. The structural analyses accompanied by the functional assays of site-directed mutants illuminate how LAT1 dynamically interacts with transportable and non-transportable compounds, including JPH203.

## Results

### LAT1 in lipid nanodiscs

We hypothesized that lipid environment is the key to investigating system L transport and inhibition. We purified LAT1–CD98hc and reconstituted it into nanodiscs (Fig. S2a), with a phospholipid mixture supplemented with cholesterol, which has been shown to be important for transport activity (*18*). To add fiducial markers for single-particle analysis, we generated mouse monoclonal antibodies and screened structure-specific binders by ELISA, FACS and negative-stain electron microscopy. The Fab fragment from clone 170 (Fab170) was found to bind to CD98hc (Fig. S2b), similar to a previously characterized antibody MEM-108 (*19*), and was used for subsequent studies. These technical improvements and sample optimization led to the structure determination of LAT1–CD98hc bound to JPH203 by cryo-EM at nominal resolution of 3.9 Å. Focused refinement on the transmembrane domain (TMD) yielded a map with better density for the ligand, with local resolutions extending to 3.6 Å (Fig. S2 and Tables S1 and S2).

### JPH203 binds to the outward-open pocket of LAT1

The structure of LAT1 bound to JPH203 shows an outward-facing conformation with the extracellular halves of TM1 and TM6 (named TM1b and TM6a) widely open (Fig. 1c). The cryo-EM map shows a well-resolved density for JPH203, which adopts a U-shaped conformer and is stuck between the hash and the bundle domains (Fig. 1d). Its α-carboxy and α-amino groups are recognized by the unwound regions of TM1 and TM6, respectively, agreeing well with the proposed binding modes for amino acid substrates (Fig. 1d). Unexpectedly, the 5-amino-2-phenylbenzoxazol side group (Fig. 1d) is not accommodated in the previously proposed “distal pocket” surrounded by TM6 and TM10 (*19*), but instead faces TM3 in an opposite direction and is partially exposed to the extracellular solvent (Fig. 1d). The three aromatic rings align nearly parallel to each other and face a flat hydrophobic patch formed by Ile140, Ser144, Ile147, Val148 and Ile397 on TM3 and TM10 (Fig. 2a). The core dichloro-Tyr moiety of JPH203 (Fig. 1d) is sandwiched by two aromatic residues: Phe252 forms a T-shaped π–π interaction with the terminal phenyl moiety of JPH203 on the extracellular side and Tyr259 forms halogen bonding interaction with chloride atom of JPH203 (Fig. 2b). Furthermore, JPH203 introduces a kink on TM10 with its nitrogen atom wedging into the helix (Fig. 2a). Together, these interactions widen the substrate binding pocket of LAT1 and fix it in the wide, outward-open conformation.

**Figure 2.**
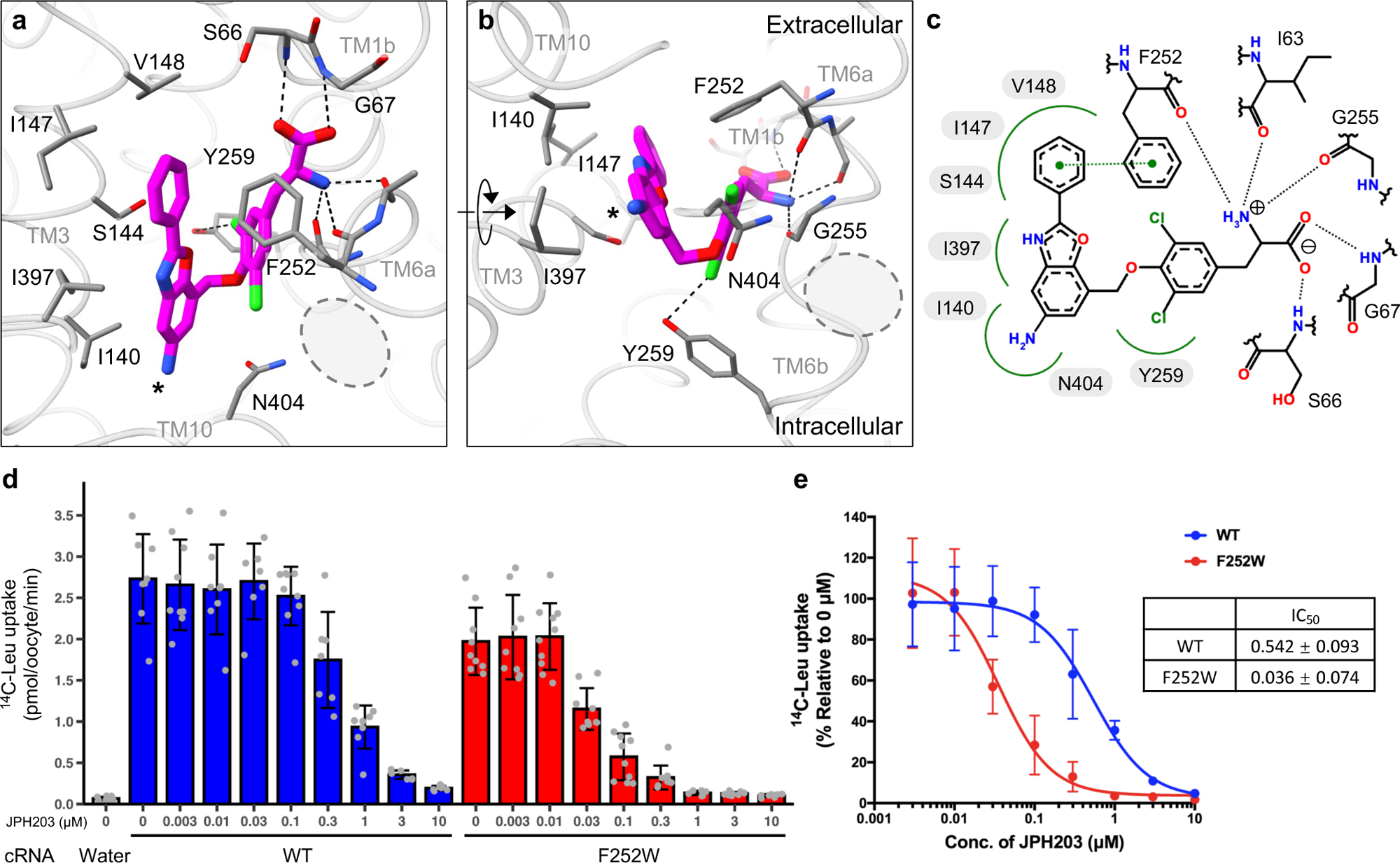
Structural basis of LAT1 inhibition by JPH203. **a,b)** Close-up views of the JPH203-binding site. Interacting residues are shown as stick models. Hydrogen bonds are depicted as dotted lines. The 5-amino group of JPH203 is marked by an asterisk. The previously predicted “distal pocket” is marked by gray dotted circles. **c)** Schematic diagram of the interactions between JPH203 and LAT1. Hydrogen bonds are depicted as black dashed lines. Hydrophobic contacts are depicted by green splines. A π–π interaction is depicted as a green dashed line connecting two green dots. **d)** Inhibition of LAT1 by JPH203 at different concentrations. Uptake of ʟ-[^14^C]Leu into *Xenopus* oocytes expressing co-expressing CD98hc and wild-type or F252W LAT1 was measured in the presence of JPH203 at indicated concentrations. As a negative control, water was injected instead of cRNAs. Data are mean ± SD and each data point represents a single oocyte (n = 7–10). **e)** Concentration-dependent inhibition curves by JPH203 for wild-type or F252W LAT1, calculated using the same data as shown in panel **d**.

JPH203 is known for its outstanding selectivity for LAT1 over LAT2 (*11*). To analyze how selectivity is conferred, we compared amino acid sequences of LAT1 and LAT2 around the JPH203-interacting residues and found key differences (Fig. 2c). For instance, Ser144 is substituted to Asn in LAT2, which may hinder the accommodation of the side group, and Phe400 to Val, which may weaken hydrophobic interactions. To test how these residues influence inhibitor sensitivity, we generated single or double variants of LAT1 and evaluated their inhibition by JPH203 using radioactive L-Leu uptake assays with *Xenopus* oocytes (Fig. S4a). Among several variants tested, the double variant F400V/S144N, which retained ∼20% of the maximal L-Leu transport activity of the wild-type, altered the sensitivity to JPH203, with almost negligible inhibition even at 1 μM, suggesting that one of these residues may be important for JPH203 selectivity (Fig. S4b). Although a single variant S144N abolished L-Leu transport and thus its inhibition could not be evaluated (Fig. S4a), the variant F400V retained the activity and the JPH203 sensitivity at around 30% (Fig. S4a,b), suggesting that Ser144 is a key residue for JPH203 selectivity. To further probe structural determinants of JPH203 recognition, we examined other amino acid substitutions (Fig. S4c). Surprisingly, F252W showed stronger inhibition by JPH203 than the wild-type, with IC_50_ value dropping from 542 nM to 36 nM (Fig. 2d,e). This may be explained by enhanced aromatic interaction between the introduced tryptophan and the terminal phenyl moiety of JPH203, consistent with the observation that adding a methoxy group to the terminal ring of JPH203 improves its affinity (*20*), presumably by enhancing the aromaticity.

We next compared the JPH203-bound LAT1 structure with those bound to JX-075, JX-078 and JX-119 (*17*). These compounds are bound to the same site in the pocket, but their detailed binding poses and structural effects on LAT1 are different (Fig. S3a–c). The core 2-amino-l,2,3,4-tetrahydro-2-naphthoic acid moiety of JX inhibitors is positioned deeper in the pocket and closer to TM3, by about 2 Å as compared to the equivalent moieties of JPH203 (Fig. S3a). In addition, the gating bundle (TM1b and TM6a) adopts a more closed conformation for the JX series than that for JPH203 (Fig. S3a). Among the three JX structures reported, only JX-119 adopts a U-shaped conformation akin to JPH203, but the blurred ligand density suggests that its terminal moiety is flexible and lacks the critical T-shaped π-π stacking with Phe252 that was observed in JPH203 (Fig. S3b,c). Interestingly, the structure of LAT1 bound to diiodo-Tyr (*17*) aligns well with the core Tyr moiety of JPH203 (Fig. S3c). These compounds share two halogen atoms at 3’- and 5’-positions, which appear to contribute to the large opening angle of TM1b and TM6a (Fig. S3c). However, due to the absence of the bulky side group, in the diiodo-Tyr-bound structure TM10 adopts a straight form (*17*), rendering the substrate-binding site not as widely open as in the JPH203-bound structure (Fig. 2a). Taken together, both the bi-halogenated Tyr core and the bulky side group of JPH203 contribute to the maximal opening of TM1b, TM6a and TM10 to ensure the wide outward-open conformation of LAT1 observed here.

### Phe and Melphalan show different degrees of interactions within the pocket

Given the successful structure determination of LAT1 bound to JPH203 in nanodiscs, we next determined the structures of LAT1 bound to a physiological substrate L-Phe and a slow substrate melphalan (Fig. 3a–c). We also serendipitously obtained an apo outward-open structure from the sample incubated with T_3_ (see Methods), which served as a reference for ligand density validation (Fig. S5; also see Methods). All three structures adopt a similar outward-facing conformation, in which TM1b and TM6a are widely open with slightly different angles (Fig. 3a–d). In the apo outward-open structure, Phe252 is flipped “up”, exposing the substrate-binding pocket towards the extracellular solvent (Fig. 3d).

**Figure 3.**
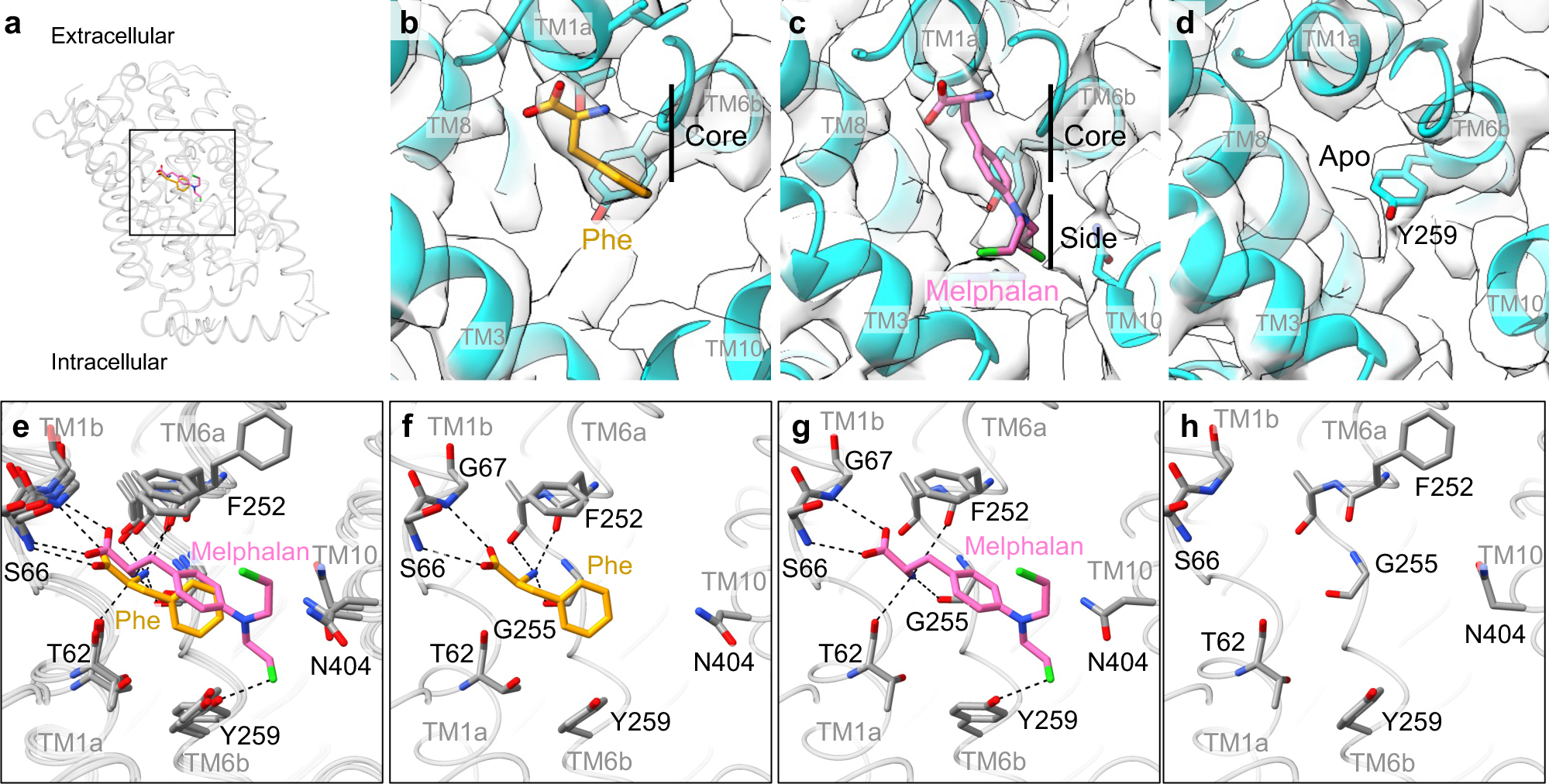
Cryo-EM structures of LAT1 bound to Phe and melphalan or in apo in the outward-facing conformation. **a)** Overlay of the Phe- and melphalan-bound LAT1, with the ligands displayed as stick models. The square indicates the region zoomed up in panels **e**–**h**. **b,f)** Structure of Phe-bound LAT1 in the outward-open conformation. **c,g)** Structure of melphalan-bound LAT1 in the outward-open conformation. **d,h)** Structure of apo LAT1 in the outward-open conformation. **e)** Superposition of the Phe-bound, melphalan-bound and apo LAT1 structures.

In the L-Phe- and melphalan-bound structures, the binding poses of the ligands agree well with previous predictions (*21*) and that of JPH203, where the substrate carboxy and amino groups are recognized by the exposed main chain atoms of TM1 and TM6, respectively (Fig. 3f,g). The phenyl ring of L-Phe faces Gly255 and forms van der Waals interactions with its Cβ atom (Fig. 3f), consistent with our previous finding that Gly255 is important for recognition of larger amino acids (*19*). Melphalan shows similar interactions in the core, but the additional bis-(2-chloroethyl)amino side group is placed in the space surrounded by TM3, TM6a and TM10 to form further interactions (Fig. 4g). Although the moderate local resolution of the map (∼ 3.9 Å) obscures precise positioning of individual atoms (Fig. 3c and Fig. S5), each of the two chloroethyl moieties appears to point upwards and downwards, with terminal chloride atoms within halogen-bonding distances to Tyr259 and Asn404 (Fig. 3f). These additional interactions would add steric hindrance to TM3, TM6b and TM10 and restrict the conformational change, which could explain its slow transport rate across the blood-brain barrier (*22*) and its inhibitory action on amino acid transport (*4*).

**Figure 4.**
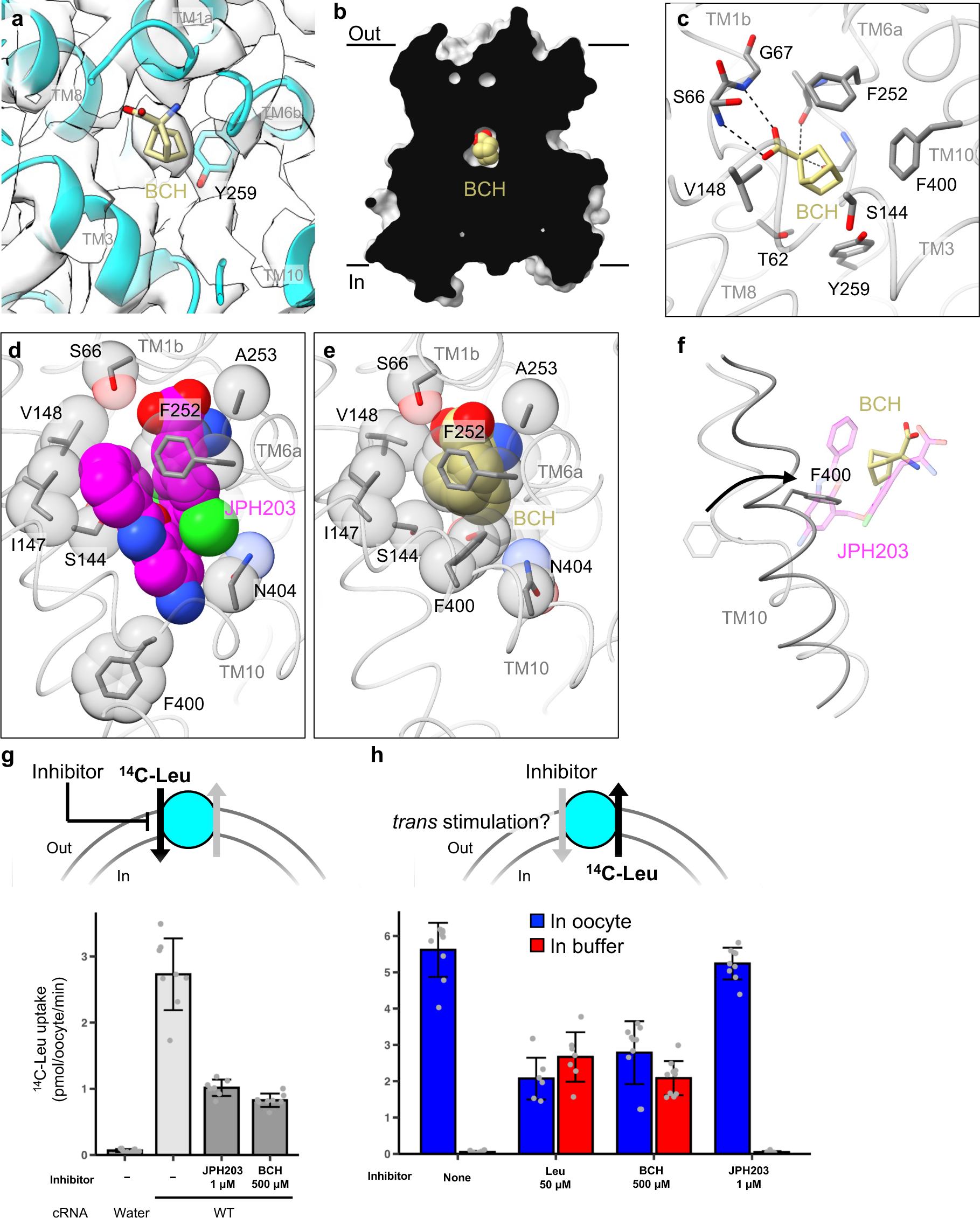
LAT1 bound to BCH in the occluded conformation. **a)** Close-up view of BCH modelled into the cryo-EM map. **b)** Cut-away surface representation of LAT1, showing that BCH is occluded from both sides of the membrane. **c)** Interaction of BCH with the surrounding residues. Dotted lines depict hydrogen bonds. **d,e)** Comparison of JPH203- and BCH-binding sites of LAT1. The ligands and important residues are shown as sticks and spheres. **f)** Rotation of Phe400 in the JPH203-bound (transparent) to BCH-bound (opaque) structures. **g)** *cis* inhibition assay. Uptake of ʟ-[^14^C]Leu into *Xenopus* oocytes expressing wild-type LAT1 and CD98hc was measured in the presence or absence of inhibitors in the external buffer solution. As a control, water was injected instead of cRNAs. Data are mean ± SD and each data point represents a single oocyte (n = 6–10). **h)** *trans* stimulation assay. Efflux of pre-loaded ʟ-[^14^C]Leu from *Xenopus* oocytes was measured in the presence or absence of indicated compounds in the external buffer solution. In the small inset, the radioactivity in the buffer was expressed as a percentage to the total radioactivity (buffer + oocytes). Data are mean ± SD and each data point represents a single oocyte (n = 7–10).

Alkylating agents are known to be transported at different rates by LAT1, acting sometimes as strong inhibitors, depending on their core structures and the positions of the mustard moiety. For example, phenylglycine-mustard (PGA) is transported at a higher rate than melphalan (*23*), whereas meta-substituted phenylalanine mustard derivatives act as potent inhibitors (*24*, *25*). Our structure suggests that the shorter core structure of PGA would pose weaker steric hindrance around TM10 to enable faster transport, whereas the bulkier substitutions especially at the meta position might add severe steric hindrance, enhancing inhibitory properties.

### Non-selective system L-inhibitor BCH induces an occluded state

We next determined the structure of LAT1 bound to a “classical” system L inhibitor BCH at 3.7 Å resolution (Fig. 4a,b). Intriguingly, unlike all the other inhibitor-bound structures, the BCH-bound LAT1 is captured in an occluded state, a previously unseen conformation where the ligand is fully sealed from both sides of the membrane (Fig. 4c). We note that this conformation considerably differs from the previous “BCH-bound”, inward-open structure (*16*), in which the modelled BCH shows poor density that is indistinguishable from that without any substrate (Fig. S1). BCH is bound in the canonical substrate-binding pocket, with its carboxy and amino moieties recognized by the exposed main chain atoms of TM1 and TM6 (Fig.4b). In comparison to the JPH203-bound structure (Fig. 4d), the pocket of BCH-bound LAT1 is significantly narrowed (Fig. 4e). TM1b and TM6a come closer to TM3 so the hydrophobic norbornane moiety of BCH contacts Ser144, Ile147 and Val148, and Phe252 sits on top of BCH to completely occlude it from the extracellular solvent (Fig. 4d,e). In addition, TM10 is twisted by about 70° to form a straight helix (Fig. 4f), bringing Phe400 in contact with the side face of the norbornane moiety. These collective structural changes collapse the space that was occupied by the 5-amino-2-phenylbenzoxazol moiety of JPH203, and the pocket is just large enough to accommodate BCH (Fig. 4c).

### JPH203 blocks but BCH facilitates transporter turnover

In the “alternating-access” scheme of membrane transporters (*26*), an occluded state represents the key intermediate that connects the outward- and inward-open states to enable substrate translocation (*27*). Our observation that BCH induces such an occluded state suggests that BCH may facilitate state transitions and thus enhance the transporter turnover. To test this, we performed transport assays using *Xenopus* oocytes expressing LAT1. In *cis* inhibition assays, in which the inhibitor is present in the external solution, BCH inhibited the uptake of the radioactive L-Leu into oocytes (Fig. 4g). This action was similar to that of JPH203, although BCH needed much higher concentrations to achieve similar level of inhibition (Fig. 4g). We next performed *trans* stimulation assays, by first pre-loading radioactive L-Leu into oocytes and then monitoring of the efflux of radioactivity upon application of compounds of interest in the external solution (Fig. 4h). The result showed that extracellular BCH indeed facilitated L-Leu efflux from the cell to the external solution (Fig. 4h). The total efflux was comparable to that stimulated by a physiological substrate L-Leu (Fig. 4h). By contrast, JPH203 did not facilitate efflux (Fig. 4h,i), consistent with its blocking action from the extracellular side (Fig. 1c). These observations demonstrate that BCH is a transportable substrate of LAT1, which acts as a competitive inhibitor when present with substrates on the same side of the membrane. This action is in stark contrast with JPH203, which, as shown above, is a *bona fide* blocker acting on the extracellular side of LAT1.

### Structural rearrangements in the transport mechanism

During image processing, we found that cryo-EM data of LAT1 incubated with some of the ligands contained subpopulation of particles representing the inward-open structures (Fig. S2g,h,i,j). In all of these, we solely observed an empty substrate-binding site, suggesting that the inward-open state is a low-affinity state that prefers ligand release. By combining all these particles, we obtained the apo inward-open structure at 3.6 Å resolution (Fig. S2k), which completed all major conformations of LAT1 and enabled detailed investigation of structural rearrangements during substrate transport (Fig. 5).

**Figure 5.**
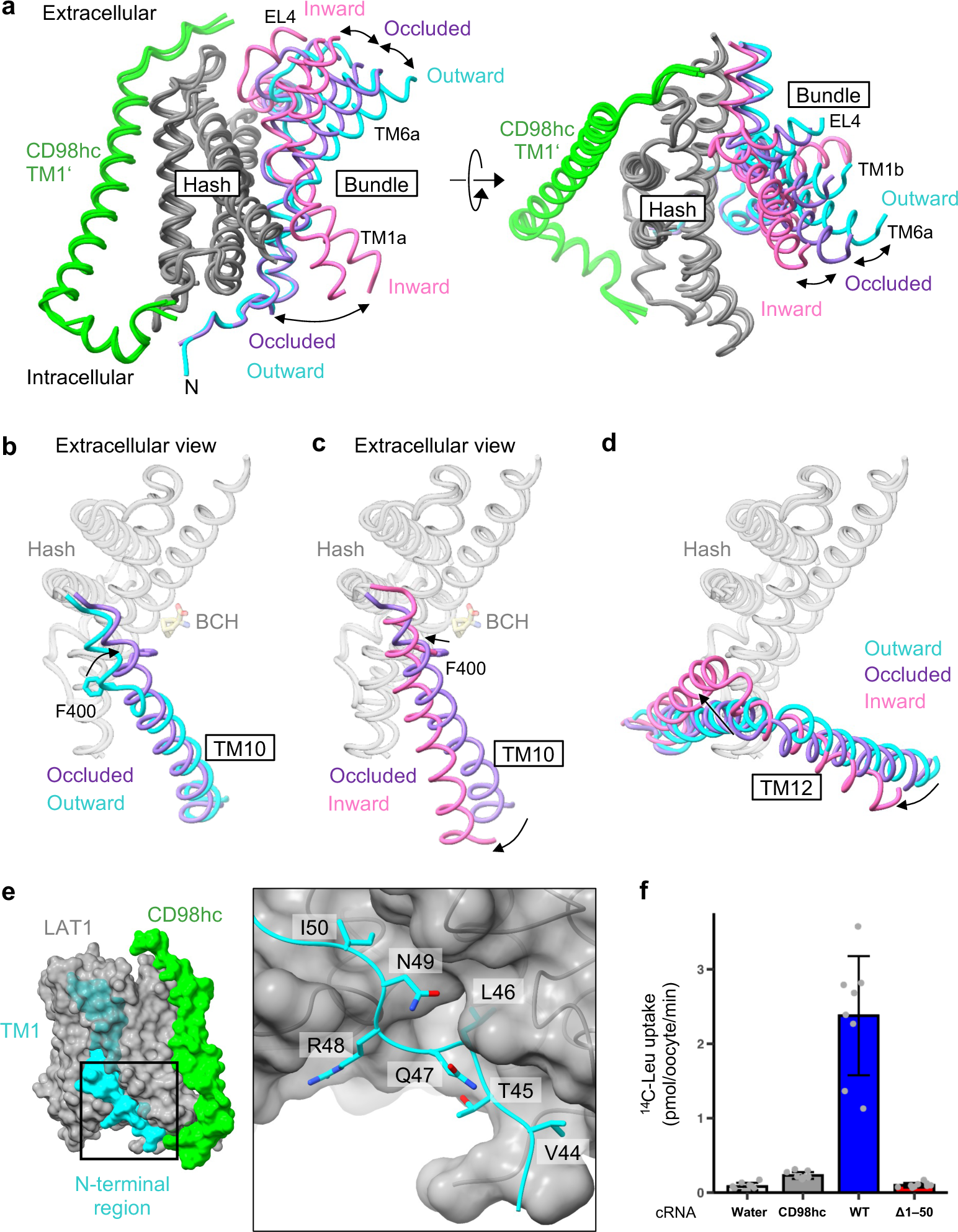
Structural rearrangements of LAT1 for substrate transport. **a)** Superposition of three major conformational states of LAT1 observed in the study, depicting the ‘rocking-bundle’ movements relevant to substrate translocation. The hash domain of LAT1 is colored gray and TM1’ of CD98hc is green. The bundle domain is colored differently for three conformations: inward-open (pink), occluded (purple) and outward-open (cyan). **b,c)** Superposition of TM10 in the three conformations. **d)** Superposition of TM12 in the three conformations. **e)** Close-up view of an N-terminal region of LAT1 in the outward-open state. **f)** Uptake of ʟ-[^14^C]Leu into *Xenopus* oocytes expressing CD98hc and wild-type or Δ1–50 LAT1. Data are mean ± SD and each data point represents a single oocyte (n = 7–10). **g)** Superposition of the whole LAT1–CD98hc complex in the three conformations.

Major structural changes occur in the gating bundle, in agreement with other well-studied LeuT-fold transporters (Fig. 5a) (*26*). In the outward-open to occluded state transition, TM6a and TM1b incline towards the hash domain by about 18° (Fig. 5a), and Phe252 sits atop the substrate, sealing the extracellular vestibule (Fig. 4e). TM10 undergoes helix rearrangements that include the rotation of about 160°, bringing Phe400 into direct hydrophobic contact with the substrate (Fig. 5b). Accompanied by these, EL4 moves closer to the CD98hc ectodomain. In the occluded to inward-facing state transition, TM6a and TM1b undergo further inward rotation towards the hash domain, accompanied by the movements of EL4, which now meets EL2 and CD98hc (Fig. 5a). TM10 undergoes a further shift of about 3 Å, to tighten the extracellular gate. Upon adopting the inward-open state, TM1a and TM6b swing open, creating an intracellular vestibule that would allow the release of substrates inside the cell (Fig. 5a,c). The binding of a counter-substrate from inside the cell would trigger the reverse process to complete an antiporter turnover.

We found that the N-terminal residues 44–50, which are disordered in the inward-facing conformations (*16*, *19*), form a defined structure in the outward-open and occluded conformations (Fig. 5e). This region forms multiple hydrophilic and hydrophobic interactions at the interface of the hash and bundle domains to strengthen the cytoplasmic gate. To test if this region plays a role in substrate transport, we generated an N-terminal truncation variant of LAT1, designated Δ1-50. This variant retained cell surface expression in oocytes (Fig. S6b,c), but showed no transport activities (Fig. 5f), demonstrating the importance of these residues for the catalytic mechanism. We also tested several other variants of this region but observed varying degrees of cell-surface localizations, which hampered identification of critical residues for substrate translocation and protein trafficking.

Notably, we observed the numerous lipid densities surrounding the TMD of LAT1– CD98hc (Fig. S7). Of these, one cholesterol bound in the cleft between TM9 and TM12 shows prominent density only in the inward-open state and gets blurred towards the occluded and outward-facing states, suggesting its conformation-specific binding (Fig. S7). Indeed, TM12 moves drastically during the transporter’s conformational change (Fig. 5d) and generates a cleft suitable for cholesterol binding only in the inward-facing state (Fig. S7c). This transient binding of cholesterol to the TM9-TM12 cleft may explain why cholesterol is needed to enhance the activity of LAT1 *in vitro* (*18*).

### Discussions

In this study, by combining lipid nanodisc and cryo-EM, we have determined the structures of LAT1 in three different conformations and revealed the mechanisms by which it recognizes physiological substrates and therapeutic compounds. Notably, JPH203 is the sole LAT1 inhibitor currently in human clinical trials for cancer treatment (*14*, *15*). Our structure provides the first atomic insight into its mechanism of action. While BCH, another system L inhibitor, has long been used to evaluate the roles of system L in various biological preparations (*28*, *29*), under the assumption that its action is solely to inhibit amino acid uptake, our structural and functional analyses have shown that it serves as a genuine substrate that can stimulate transporter turnover. This would call for careful re-evaluation of *in vivo* and *in vitro* studies in which BCH was applied extracellularly, as it may not only inhibit the uptake of neutral amino acids but also enhance their efflux. Our study has also revealed the recognition mechanism of melphalan, a chemotherapeutic agent that is known to be slowly transported by system L (*4*), shedding light on the mechanisms by which various bioactive amino acid derivatives interact with LAT1.

Previous pharmacophore modelling suggested that the substrate-binding pocket of LAT1 has a large hydrophobic “free” space capable of accommodating the hydrophobic side groups of inhibitors or prodrugs (*4*, *30*). Our structural findings demonstrate that this long-predicted hydrophobic space is not a rigid, pre-defined pocket. Instead, it comprises highly mobile structural elements, namely TM6, TM10 and part of TM3, which dynamically adapt their conformations to fit the ligands of varying shapes. For instance, in the case of JPH203, this mechanism is exploited to block the transporter, where the amino-phenylbenzoxazol moiety wedges into TM10 to expand the hydrophobic space and arrest the conformational change (Fig. 6). In contrast, BCH shows an opposite action, where the norbornane moiety attracts TM10 to narrow the hydrophobic pocket and facilitate the conformational change (Fig. 6). Notably, the occluded pocket observed for BCH is too narrow to accomodate melphalan. This aligns with the observed slower transport rate of melphalan and other large amino acid derivatives such as T_3_. We propose that the transport process of these large amino acid derivatives may involve a “loose” occluded state, where the substrate is not as tightly confined by surrounding residues as it is for BCH, yet it could still be translocated across the membrane by a series of conformational changes. Another possibility is that the previously proposed “distal pocket” plays a role in the transport of such larger substrates.

**Figure 6.**
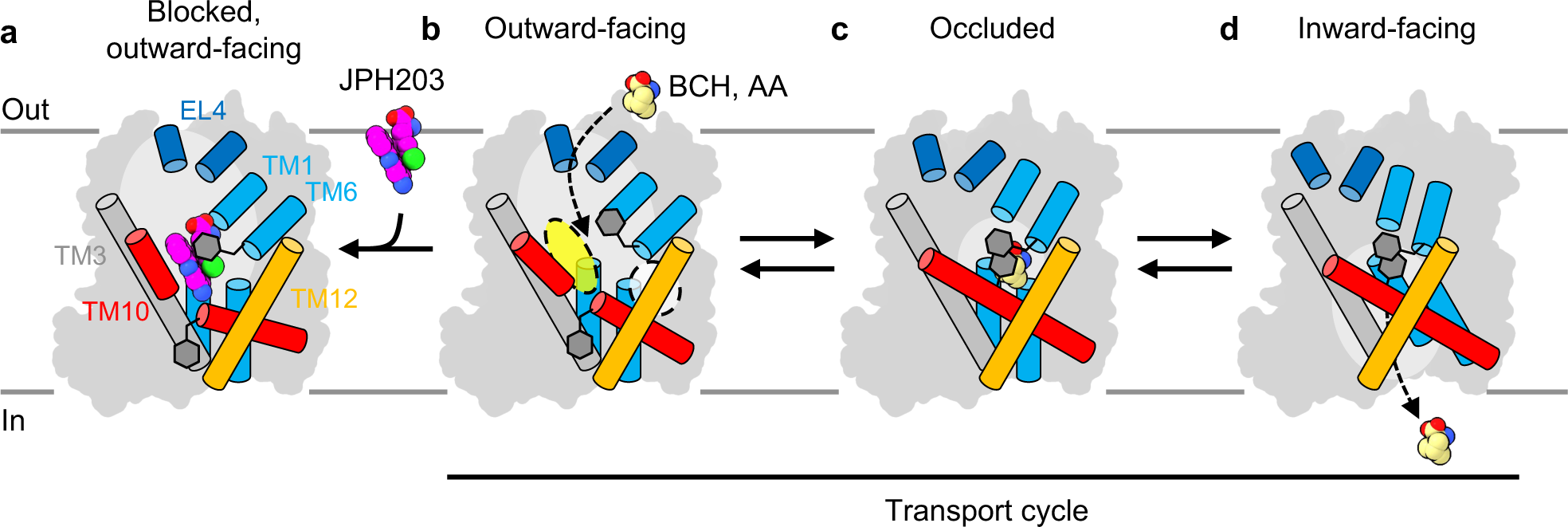
Transport cycle of LAT1 and inhibition mechanisms. **a–d)** Schematic representation of amino acid transport by LAT1 and its inhibition by JPH203 and BCH. All four panels represent experimental structures reported in the study. In the outward-facing state (**a**,**b**), LAT1 can accept substrates from extracellular solvent, with mobile Phe252 on TM6 facilitating access (**b**). The white transparent circle depicts the distal pocket predicted previously, and the yellow circle the hydrophobic space found in this study. When JPH203 binds (**a**), TM10 is bent, Phe252 closes, and TM1b and TM6a are pushed open, locking the transporter in the outward-facing state and blocking substrate access. Binding of BCH induces the occluded conformation (**c**), characterized by the rotation of Phe400 on TM10 toward the pocket. Eventually LAT1 transitions to the inward-facing state (**d**), which has low affinity for the substrate and releases it into the cell.

In conclusion, the structures of LAT1 bound with substrates and inhibitors provide a basis for understanding its amino acid transport and inhibition mechanisms, and establish a blueprint for the better design of transportable or non-transportable drugs targeting system L. Additionally, the nanodisc system employed here holds promise for further structural and biochemical characterization of LAT1.

## Methods

### Protein expression and purification

Human LAT1–CD98hc was expressed and purified essentially as described (*19*). To improve structural integrity of the protein, we modified purification protocols in several ways. First, we used high-expression batches of HEK293 cells, by using concentrated baculoviruses for maximal infection efficiency of two components (*31*). Second, we skipped a FLAG-tag purification step and used only the GFP-nanotrap for affinity purification. Third, the wash buffer for GFP-nanotrap was supplemented with phospholipid mixture (0.0025% POPC, 0.0025% POPG (w/w)). Finally, gel filtration in the presence of detergent was skipped, and the GFP-nanotrap eluate was directly concentrated and used for nanodisc reconstitution. For nanodisc reconstitution, purified protein was mixed with POPC:POPG:cholesterol (2:2:1 (w/w)) and MSP2N2 at a 1:5:100 molar ratio and incubated overnight at 4 degrees, with the step-wise addition of Bio-Beads SM-2 as described (*32*). The reconstituted complex was further purified by size-exclusion chromatography (SEC) on a Superose 6 Increase 3.2/300 column with the SEC buffer (20 mM Tris-HCl, pH 9.0, 150 mM NaCl).

### Monoclonal Antibody generation and selection

The animal experiments conformed to the guidelines outlined in the Guide for the Care and Use of Laboratory Animals of Japan and were approved by the Chiba University Animal Care Committee (approval number. 30-174), and performed according to ARRIVE guidelines. 4-week-old female MRL/MpJJmsSlc-*lpr/lpr* mice were purchased from Nihon SLC (Shizuoka, Japan). The temperature of the animal breeding room was controlled at 20–25°C, and the lighting was alternated between day and night at 12 h/12 h. Mice were immunized with liposomes made of purified LAT1–CD98hc and Egg PC (Avanti Polar Lipids, Inc., Birmingham, AL.). After several rounds of immunization, splenocytes from the immunized mice were fused with P3U1 myeloma cells and generated hybridomas. Conformational and extracellular-domain recognizing antibodies produced by hybridomas were screened by using liposome-ELISA, antigen-denatured-ELISA (*33*) and flow cytometry. IgGs from established clones were purified by Protein G column (Cytiva, Inc., Marlborough, MA.) from culture supernatants and digested into Fab fragments by Papain (Nacalai tesque, Inc., Kyoto, Japan), and further purified on Protein A column (Cytiva). A total of 20 clones were assessed for their binding sites on LAT1–CD98hc using negative-stain microscopy. The Fab fragment from clone 170 (Fab170), which showed rigid binding on the extracellular epitope of CD98hc, similar to a previously characterized MEM-108 (*19*), was used for the structural studies.

### Cryo-EM sample preparation

Nanodisc-reconstituted LAT1–CD98hc was complexed with Fab170, subjected gel filtration and concentrated to about 15 mg/ml for cryo-EM sample preparation. 3 μl of sample was applied to glow-discharged Quantifoil or C-flat holey carbon grids (copper, 400 mesh, 1.2/1.3 hole size) and blotted for 3–4 seconds using Vitrobot Mark I. To improve particle distribution, 1.5 mM fluorinated Fos-Choline-8 was added immediately before sample application. For JPH203, the inhibitor was added from a 2 mM stock solution in DMSO to a final inhibitor concentration of 20 μM and the mixture was further incubated at 37°C for 10 min to ensure full binding. For L-Phe (final 5 mM), melphalan (final 400 μM), BCH (final 30 mM) and T_3_ (final 50 μM), the ligands were added to the sample approximately 1 hour before vitrification and kept on ice. For T_3_, we also tried longer incubation, by including T_3_ before the final gel filtration step and all subsequent steps, but neither of the strategies yielded T_3_-bound LAT1 structures.

### Cryo-EM data acquisition, processing and structure determination

A total of 84,295 cryo-EM movies were collected on multiple separate sessions, summarized in Fig. S2d–j. All data were collected on the same Titan Krios G3 microscope equipped with a BioQuantum K3 camera at a nominal magnification of 105 k×, which corresponds to a calibrated pixel size of 0.837 Å /pix. The camera was operated in the counted super-resolution mode with a binning factor of 2 and the energy filter slit width was set to 30 eV. The electron flux rate was 15 e^−^/pix/sec, the exposure time 2.5 sec, and the total exposure 51 e^−^/Å^2^. All movies were fractionated into 50 frames. Data were collected with EPU software with the aberration-free image shift method. Data quality was monitored with cryoSPARC Live v2.15 (*34*).

All cryo-EM data processing was performed in RELION 3.1 (*35*, *36*). Movies were motion-corrected in MotionCor2 (*37*) with 5 × 5 patches and the contrast transfer function (CTF) was estimated in CTFFIND4 (*38*). Particles were picked with a Laplacian-of-Gaussian picker and Topaz (*39*) and extracted with down-sampling to a pixel size of 3.45 Å. All good particles after rounds of 2D and 3D classifications were combined, duplicates removed, and the non-redundant particles were extracted with a final pixel size of 1.5345 Å. Further 3D classification showed that the L-Phe, JPH203, T_3_ (long incubation) and no-substrate datasets contained only one conformation, whereas the melphalan, BCH or T_3_ (short incubation) datasets contained multiple conformations. Focused classification showed that these datasets contained the apo inward-open structures to a varying ratio, which were further classified and combined to yield a single reconstruction. After combining the particles of the same conformation, Bayesian polishing (*40*) was performed with trained parameters.

Even after separating the discrete conformational changes, the continuous structural flexibility of each component (LAT1, CD98hc, Fab and the surrounding nanodisc) hampered high-resolution reconstruction for the region of interest. To overcome this, we resorted to multibody refinement (*41*). Details of the refinement parameters and particle selection strategies are described elsewhere (manuscript in preparation). Briefly, three bodies were defined, designated as the “core”, “Fab” and “TMD” (Fig. S2l). The “core” mask includes LAT1–CD98hc excluding the nanodisc and Fab170, the “Fab” mask includes only Fab170, and the “TMD” mask includes LAT1, TM1’ of CD98hc and the entire nanodisc. Particles were selected through repeated “refine, subtract and classify” cycles, and the final reconstructions of the “TMD” body from multibody refinements were used for model building of the TMD for all datasets. The models for the whole complex (LAT1–CD98hc–Fab170) were built only for the JPH203-bound, BCH-bound and inward-open apo states by using the consensus map as representatives of the outward-facing, occluded and inward-facing conformations, respectively.

A published inward-open structure (PDB ID: 6IRS) was first rigid-body fitted into the maps in ChimeraX 1.0 (*42*). Subsequent model building was performed in COOT 0.9 (*43*) with ligand restraints generated in AceDRG (*44*). Refinement was performed in Servalcat (*45*) using two unfiltered half-maps. The BCH compound (Sigma) contains exo- and endo-carboxy isomers with an unknown ratio. We chose an isomer with an exo-carboxylic group for modelling, based on a previous NMR study of BCH from the same supplier (*46*). Data collection and refinement statistics are shown in Tables S1 and S2.

### Cryo-EM ligand validation

For ligand validation in cryo-EM maps, we utilized map validation tools in REFMAC5 (*47*). First, we used Servalcat (*45*) to calculate “omit” Fo – Fc difference maps from the two unfiltered half maps and the ligand-omitted models for each ligand-bound state. The Fo – Fc maps clearly revealed the presence of the ligands (Fig. S5). Second, we used EMDA (*48*) to calculate Fo – Fo maps between the ligand-bound and apo states. Maps were first fitted to each other using FSC-based auto-resolution threshold and providing a “TMD” mask. Then, the apo map was subtracted from each ligand-bound map. This analysis revealed clear densities for all ligands, JPH203, L-Phe, melphalan and BCH, fitting well the models (Fig. S5).

### Transport measurements and expression analysis using X. laevis oocytes

Functional and expression analyses of LAT1 using *X. laevis* oocytes were conducted as described previously unless otherwise specifically denoted (*19*). LAT1 mutants with amino acid substitutions or an *N*-terminal truncation (Δ1-50) were constructed by whole-plasmid PCR using PrimeSTAR MAX DNA polymerase (Takara). The corresponding codons were altered as follows for amino acid substitution: S144N (AAT), F252Y (TAT), F252W (TGG), Y259F (TTC), F394Y (TAT), F400V (GTG), F400I (ATT), F400L (TTA), and F400W (TGG).

Uptake experiments of LAT1 in *X. laevis* oocytes were performed in Na^+^-free ND96 buffer containing 50 μM of L-[^14^C]Leu (3.3CCiCmol^−1^, Moravek) for 30 min at room temperature. The indicated concentrations of JPH203 or BCH were added to the buffer when specified. After the lysis of oocytes with 10% (v/v) SDS, the radioactivity was determined using a β-scintillation counter (LSC-3100, Aloka, Tokyo, Japan). In efflux experiments, oocytes were preinjected with 50 nL of 100 μM L-[^14^C]Leu (3.3 nCi/ oocyte). After an extensive wash with ice-cold Na^+^-free ND96 buffer, the oocytes were incubated in the buffer with or without the indicated concentrations of JPH203, BCH, or L-Leu for 15 min at room temperature to induce efflux of preloaded L-[^14^C]Leu. Then the radioactivity in the buffer and the remaining radioactivity in the oocytes were separately counted. L-[^14^C]Leu efflux was expressed as a percentage of total radioactivity (the radioactivity in the buffer divided by the sum of the radioactivity of the buffer and the remaining radioactivity in oocytes).

Detection of LAT1 by immunoblotting in the total membranes of *X. laevis* oocytes was performed with anti-LAT1 (1:2,000, KE026; TransGenic) and peroxidase goat anti-rabbit IgG (1:10,000, Jackson ImmunoResearch). Immunofluorescence detection of LAT1 in paraffin sections of *X. laevis* oocytes was performed with anti-LAT1 (1:500, J-Pharma) and Alexa Fluor 488-conjugated anti-mouse IgG (A21202, 1:1000, Invitrogen). Images were acquired using a fluorescence microscope (BZ-9000, Keyence) equipped with a ×100 objective lens (CFI Plan Apo λ, numerical aperture 1.40, Nikon). Reproducibility of the results was confirmed by independent experiments using different batches of oocytes.

### Materials for radioactive assays

^14^C-leucine (338 mCi/mmol) was purchased from Moravek Biochemicals (Brea, CA). Standard amino acids and 2-aminobicyclo[2.2.1]heptane-2-carboxylic acid (BCH) were purchased from Sigma-Aldrich (St Louis, MO). JPH203 ((S)-2-amino-3-(4-((5-amino-2-phenylbenzo [d]oxazol-7-yl)methoxy)-3,5-dichlorophenyl) propanoic acid, CAS No. 1037592–40-7) (2HCl salt; purity >C99%) was provided by J-Pharma Co., Ltd (Tokyo, Japan). Unless otherwise stated, other chemicals were purchased from Wako Pure Chemical Industries (Osaka, Japan).

## Data availability

The atomic coordinates have been deposited to Protein Data Bank under accession numbers 8KDD, 8KDF, 8KDG, 8KDH, 8KDI, 8KDJ, 8KDN, 8KDO and 8KDP. Cryo-EM maps have been deposited to Electron Microscopy Data Bank under accession numbers EMD-37132, EMD-37134, EMD-37135, EMD-37136, EMD-37137, EMD-37138, EMD-37140, EMD-37141 and EMD-37142. All other data will be available upon request.

## Acknowledgments

We thank Werner Kühlbrandt for providing research infrastructure and discussions; Deryck J. Mills for microscope management and training; Susann Kaltwasser, Simone Prinz, Mark Linder and Sonja Welsch in the Central Electron Microscopy Facility of the Max Planck Institute of Biophysics for technical assistance in electron microscopy; Sabine Häder, Christina Kunz and Heidi Betz for assistance in laboratory experiments; Juan Castillo, Özkan Yildiz, the Central IT team and the Max Planck Computing and Data Facility for maintaining the computational infrastructure; and J-Pharma for providing JPH203. This work was supported by the Max Planck Society, Kazato Research Foundation, JSPS KAKENHI (21K15031), The Uehara Memorial Foundation, and Basis for Supporting Innovative Drug Discovery and Life Science Research (BINDS) from AMED (JP23ama121013) to Y.L.; The Medical Research Council, as part of UK Research and Innovation (MC_UP_A025_1012) to K.Y. and G.M. Y.L. was supported by Toyobo Biotechnology Foundation Fellowship and Human Frontier Science Program Long-Term Fellowship.

## Author contributions

Y.L., O.N. and Y.K. initiated the project. Y.L. performed EM sample preparation, data collection and structure determination. C.J. and R.O. prepared mutants and performed functional assays. R.O. and M.X. performed immunostaining and western blotting. S.O. and T.M. generated monoclonal antibodies and performed antibody screening. R.W. helped with EMDA overlay and Fo – Fo difference map calculation. K.Y. assisted with map and model validation using Servalcat. Y.L. and Y.K. wrote the manuscript, with contributions from all co-authors.

## References

1. H. Christensen, Role of amino acid transport and countertransport in nutrition and metabolism. Physiol Rev 70, 43–77 (1990).

2. F. Verrey, System L: heteromeric exchangers of large, neutral amino acids involved in directional transport. Pflügers Archiv 445, 529–533 (2003).

3. Y. Kanai, H. Segawa, K. i Miyamoto, H. Uchino, E. Takeda, H. Endou, Expression cloning and characterization of a transporter for large neutral amino acids activated by the heavy chain of 4F2 antigen (CD98). J Biol Chem 273, 23629–32 (1998).

4. H. Uchino, Y. Kanai, D. Kim, M. F. Wempe, A. Chairoungdua, E. Morimoto, M. Anders, H. Endou, Transport of amino acid-related compounds mediated by L-type amino acid transporter 1 (LAT1): insights into the mechanisms of substrate recognition. Mol Pharmacol 61, 729–37 (2002).

5. E. C. Friesema, R. Docter, E. P. Moerings, F. Verrey, E. P. Krenning, G. Hennemann, T. J. Visser, Thyroid hormone transport by the heterodimeric human system L amino acid transporter. Endocrinology 142, 4339–48 (2001).

6. Y. Kanai, Amino acid transporter LAT1 (SLC7A5) as a molecular target for cancer diagnosis and therapeutics. Pharmacol Therapeut, 107964 (2021).

7. Y. Mishima, C. Honda, M. Ichihashi, H. Obara, J. Hiratsuka, H. Fukuda, H. Karashima, T. Kobayashi, K. Kanda, K. Yoshino, TREATMENT OF MALIGNANT MELANOMA BY SINGLE THERMAL NEUTRON CAPTURE THERAPY WITH MELANOMA-SEEKING 10B-COMPOUND. Lancet 334, 388–389 (1989).

8. C. Plathow, W. A. Weber, Tumor Cell Metabolism Imaging. J Nucl Med 49, 43S–63S (2008).

9. D. Dickens, S. D. Webb, S. Antonyuk, A. Giannoudis, A. Owen, S. Rädisch, S. S. Hasnain, M. Pirmohamed, Transport of gabapentin by LAT1 (SLC7A5). Biochem Pharmacol 85, 1672–1683 (2013).

10. T. Kageyama, M. Nakamura, A. Matsuo, Y. Yamasaki, Y. Takakura, M. Hashida, Y. Kanai, M. Naito, T. Tsuruo, N. Minato, S. Shimohama, The 4F2hc/LAT1 complex transports l-DOPA across the blood–brain barrier. Brain Res 879, 115–121 (2000).

11. K. Oda, N. Hosoda, H. Endo, K. Saito, K. Tsujihara, M. Yamamura, T. Sakata, N. Anzai, M. F. Wempe, Y. Kanai, H. Endou, LCType amino acid transporter 1 inhibitors inhibit tumor cell growth. Cancer Sci 101, 173–179 (2010).

12. C. Rosilio, M. Nebout, V. Imbert, E. Griessinger, Z. Neffati, J. Benadiba, T. Hagenbeek, H. Spits, J. Reverso, D. Ambrosetti, J.-F. Michiels, B. Bailly-Maitre, H. Endou, M. Wempe, J.-F. Peyron, L-type amino-acid transporter 1 (LAT1): a therapeutic target supporting growth and survival of T-cell lymphoblastic lymphoma/T-cell acute lymphoblastic leukemia. Leukemia 29, 1253–1266 (2015).

13. P. Häfliger, J. Graff, M. Rubin, A. Stooss, M. S. Dettmer, K.-H. Altmann, J. Gertsch, R.-P. Charles, The LAT1 inhibitor JPH203 reduces growth of thyroid carcinoma in a fully immunocompetent mouse model. J Exp Clin Canc Res 37, 234 (2018).

14. N. Okano, D. Naruge, K. Kawai, T. Kobayashi, F. Nagashima, H. Endou, J. Furuse, First-in-human phase I study of JPH203, an L-type amino acid transporter 1 inhibitor, in patients with advanced solid tumors. Invest New Drug 38, 1495–1506 (2020).

15. J. Furuse, M. Ikeda, M. Ueno, M. Furukawa, C. Morizane, T. Takehara, T. Nishina, A. Todaka, N. Okano, K. Hara, Y. Nakai, K. Ohkawa, T. Sasaki, K. Sugimori, N. Yokoyama, K. Yamamoto, Nanvuranlat, an L-type amino acid transporter (LAT1) inhibitor for patients with pretreated advanced refractory biliary tract cancer (BTC): Primary endpoint results of a randomized, double-blind, placebo-controlled phase 2 study. J Clin Oncol 41, 494–494 (2023).

16. R. Yan, X. Zhao, J. Lei, Q. Zhou, Structure of the human LAT1–4F2hc heteromeric amino acid transporter complex. Nature 568, 127–130 (2019).

17. R. Yan, Y. Li, J. Müller, Y. Zhang, S. Singer, L. Xia, X. Zhong, J. Gertsch, K.-H. Altmann, Q. Zhou, Mechanism of substrate transport and inhibition of the human LAT1-4F2hc amino acid transporter. Cell Discov 7, 16 (2021).

18. D. Dickens, G. N. Chiduza, G. S. Wright, M. Pirmohamed, S. V. Antonyuk, S. S. Hasnain, Modulation of LAT1 (SLC7A5) transporter activity and stability by membrane cholesterol. Sci Rep 7, 43580 (2017).

19. Y. Lee, P. Wiriyasermkul, C. Jin, L. Quan, R. Ohgaki, S. Okuda, T. Kusakizako, T. Nishizawa, K. Oda, R. Ishitani, T. Yokoyama, T. Nakane, M. Shirouzu, H. Endou, S. Nagamori, Y. Kanai, O. Nureki, Cryo-EM structure of the human L-type amino acid transporter 1 in complex with glycoprotein CD98hc. Nat Struct Mol Biol 26, 510–517 (2019).

20. J. Zaugg, X. Huang, F. Ziegler, M. Rubin, J. Graff, J. Müller, R. MoserCHässig, T. Powell, J. Gertsch, K. Altmann, C. Albrecht, Small molecule inhibitors provide insights into the relevance of LAT1 and LAT2 in maternoCfoetal amino acid transport. J. Cell. Mol. Med. 24, 12681–12693 (2020).

21. E. G. Geier, A. Schlessinger, H. Fan, J. E. Gable, J. J. Irwin, A. Sali, K. M. Giacomini, Structure-based ligand discovery for the Large-neutral Amino Acid Transporter 1, LAT-1. Proc Natl Acad Sci U S A 110, 5480–5485 (2013).

22. Q. R. Smith, Carrier-mediated transport to enhance drug delivery to brain. Int Congr Ser 1277, 63–74 (2005).

23. K. Hosoya, H. Kyoko, N. Toyooka, A. Kato, M. Orihashi, M. Tomi, M. Tachikawa, Evaluation of Amino Acid-Mustard Transport as L-Type Amino Acid Transporter 1 (LAT1)-Mediated Alkylating Agents. Biol Pharm Bulletin 31, 2126–2130 (2008).

24. D. R. Haines, R. W. Fuller, S. Ahmad, D. T. Vistica, V. E. Marquez, Selective cytotoxicity of a system L specific amino acid nitrogen mustard. J Med Chem 30, 542–547 (1987).

25. J. Matharu, J. Oki, D. R. Worthen, Q. R. Smith, P. A. Crooks, Regiospecific and conformationally restrained analogs of melphalan and dl-2-NAM-7 and their affinities for the large neutral amino acid transporter (system LAT1) of the blood–brain barrier. Bioorg Med Chem Lett 20, 3688–3691 (2010).

26. D. Drew, O. Boudker, Shared molecular mechanisms of membrane transporters. Annu Rev Biochem 85, 1–30 (2015).

27. M. Klingenberg, Ligand−Protein Interaction in Biomembrane Carriers. The Induced Transition Fit of Transport Catalysis †. Biochemistry 44, 8563–8570 (2005).

28. C. S. Kim, S.-H. Cho, H. S. Chun, S.-Y. Lee, H. Endou, Y. Kanai, D. K. Kim, BCH, an Inhibitor of System L Amino Acid Transporters, Induces Apoptosis in Cancer Cells. Biol. Pharm. Bull. 31, 1096–1100 (2008).

29. Y. Saito, T. Soga, Amino acid transporters as emerging therapeutic targets in cancer. Cancer Sci. 112, 2958–2965 (2021).

30. H. Ylikangas, L. Peura, K. Malmioja, J. Leppänen, K. Laine, A. Poso, M. Lahtela-Kakkonen, J. Rautio, Structure–activity relationship study of compounds binding to large amino acid transporter 1 (LAT1) based on pharmacophore modeling and in situ rat brain perfusion. Eur J Pharm Sci 48, 523–531 (2013).

31. C. L. Morales-Perez, C. M. Noviello, R. E. Hibbs, Manipulation of Subunit Stoichiometry in Heteromeric Membrane Proteins. Structure 24, 797–805 (2016).

32. F. Hagn, M. L. Nasr, G. Wagner, Assembly of phospholipid nanodiscs of controlled size for structural studies of membrane proteins by NMR. Nat Protoc 13, 79–98 (2018).

33. T. Hino, T. Arakawa, H. Iwanari, T. Yurugi-Kobayashi, C. Ikeda-Suno, Y. Nakada-Nakura, O. Kusano-Arai, S. Weyand, T. Shimamura, N. Nomura, A. D. Cameron, T. Kobayashi, T. Hamakubo, S. Iwata, T. Murata, G-protein-coupled receptor inactivation by an allosteric inverse-agonist antibody. Nature 482, 237–240 (2012).

34. A. Punjani, J. L. Rubinstein, D. J. Fleet, M. A. Brubaker, cryoSPARC: algorithms for rapid unsupervised cryo-EM structure determination. Nat. Methods 14, 290–296 (2017).

35. S. H. W. Scheres, RELION: Implementation of a Bayesian approach to cryo-EM structure determination. J Struct Biol 180, 519–530 (2012).

36. J. Zivanov, T. Nakane, B. O. Forsberg, D. Kimanius, W. J. Hagen, E. Lindahl, S. H. Scheres, New tools for automated high-resolution cryo-EM structure determination in RELION-3. eLife 7, e42166 (2018).

37. S. Q. Zheng, E. Palovcak, J.-P. Armache, K. A. Verba, Y. Cheng, D. A. Agard, MotionCor2: anisotropic correction of beam-induced motion for improved cryo-electron microscopy. Nat Methods 14, 331–332 (2017).

38. A. Rohou, N. Grigorieff, CTFFIND4: Fast and accurate defocus estimation from electron micrographs. J. Struct. Biol. 192, 216–221 (2015).

39. T. Bepler, A. Morin, M. Rapp, J. Brasch, L. Shapiro, A. J. Noble, B. Berger, Positive-unlabeled convolutional neural networks for particle picking in cryo-electron micrographs. Nat Methods 16, 1153–1160 (2019).

40. J. Zivanov, T. Nakane, S. H. W. Scheres, A Bayesian approach to beam-induced motion correction in cryo-EM single-particle analysis. IUCrJ 6, 5–17 (2019).

41. T. Nakane, D. Kimanius, E. Lindahl, S. H. Scheres, Characterisation of molecular motions in cryo-EM single-particle data by multi-body refinement in RELION. eLife 7, e36861 (2018).

42. T. D. Goddard, C. C. Huang, E. C. Meng, E. F. Pettersen, G. S. Couch, J. H. Morris, T. E. Ferrin, UCSF ChimeraX: Meeting modern challenges in visualization and analysis. Protein Sci 27, 14–25 (2018).

43. P. Emsley, B. Lohkamp, W. G. Scott, K. Cowtan, Features and development of Coot. Acta Crystallogr Sect D Biol Crystallogr 66, 486–501 (2010).

44. F. Long, R. A. Nicholls, P. Emsley, S. Gražulis, A. Merkys, A. Vaitkus, G. N. Murshudov, AceDRG: a stereochemical description generator for ligands. Acta Crystallogr. Sect. D: Struct. Biol. 73, 112–122 (2017).

45. K. Yamashita, C. M. Palmer, T. Burnley, G. N. Murshudov, Cryo-EM single-particle structure refinement and map calculation using Servalcat. Acta Crystallogr Sect D 77, 1282–1291 (2021).

46. S. Bröer, A. Bröer, J. T. Hansen, W. A. Bubb, V. J. Balcar, F. A. Nasrallah, B. Garner, C. Rae, Alanine metabolism, transport, and cycling in the brain. J Neurochem 102, 1758–1770 (2007).

47. A. Brown, F. Long, R. A. Nicholls, J. Toots, P. Emsley, G. Murshudov, Tools for macromolecular model building and refinement into electron cryoCmicroscopy reconstructions. Acta Crystallogr. Sect. D 71, 136–153 (2015).

48. R. Warshamanage, K. Yamashita, G. N. Murshudov, EMDA: A Python package for Electron Microscopy Data Analysis. J Struct Biol 214, 107826 (2022).

