## Supplementary material for "Structural basis of anticancer drug recognition and amino acid transport by LAT1": Figures S1-S7 and Tables S1-S2

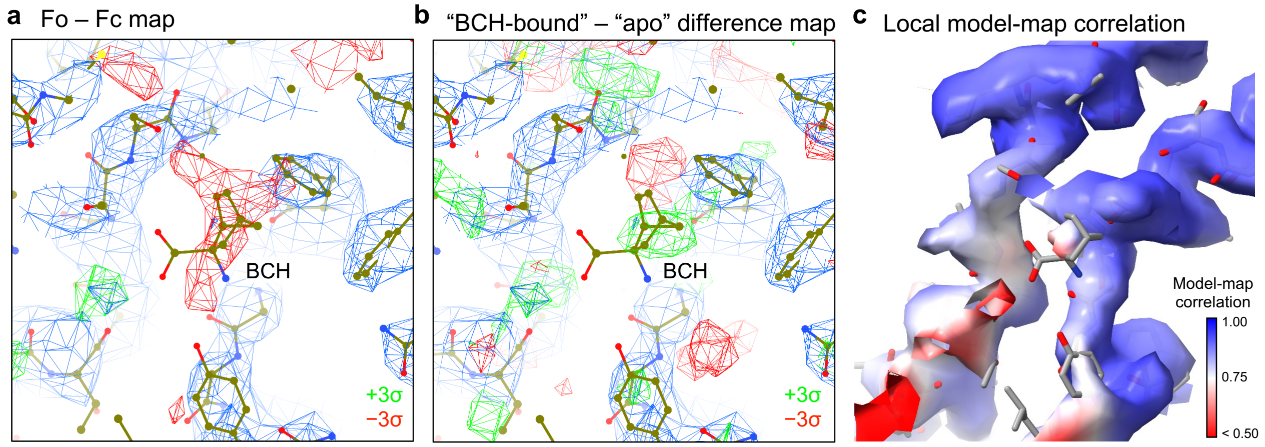


**Figure S1 | Re-analysis of a previously reported "BCH-bound" structure (PDB ID: 6IRT and EMDB ID: EMD-9722)**

**a)** Fo – Fc map calculated for the "BCH-bound" structure [Yan *et al.*, *Nature*, 2019] using the deposited PDB coordinate (6IRT) and the MRC map (EMD-9722). The blue mesh represents the original unprocessed map, and the green and red meshes represent the positive and negative Fo – Fc densities contoured at +3σ and −3σ (normalized within the mask), respectively. Since the original B-factors were set to 20.00 for all atoms of BCH in the deposited coordinates, the difference map was calculated after 10 cycles of coordinate and ADP refinement. After the refinement, the average B-factor of BCH has increased to 192.7 (ranging from 156.8 to 216.3), well above those of the surrounding atoms (eight residues within 4 Å of BCH), ranging from 25.0 to 115.3. The negative densities overlapping with BCH indicates no binding or low occupancy of the ligand.

**b)** A difference map between the "BCH-bound" (EMD-9722) and the "apo" maps (EMD-9721). The positive (green) and negative (red) densities were contoured at +3σ and −3σ (normalized within the mask), respectively. There is only a weak positive density partially overlapping with BCH, indicating no binding or low occupancy of the ligand.

**c)** Local model-map correlation calculated for the "BCH-bound" structure after refinement (6IRT and EMD-9722). The region around the modelled BCH shows poor correlation, probably due to the forced placement of the BCH in a weak density.

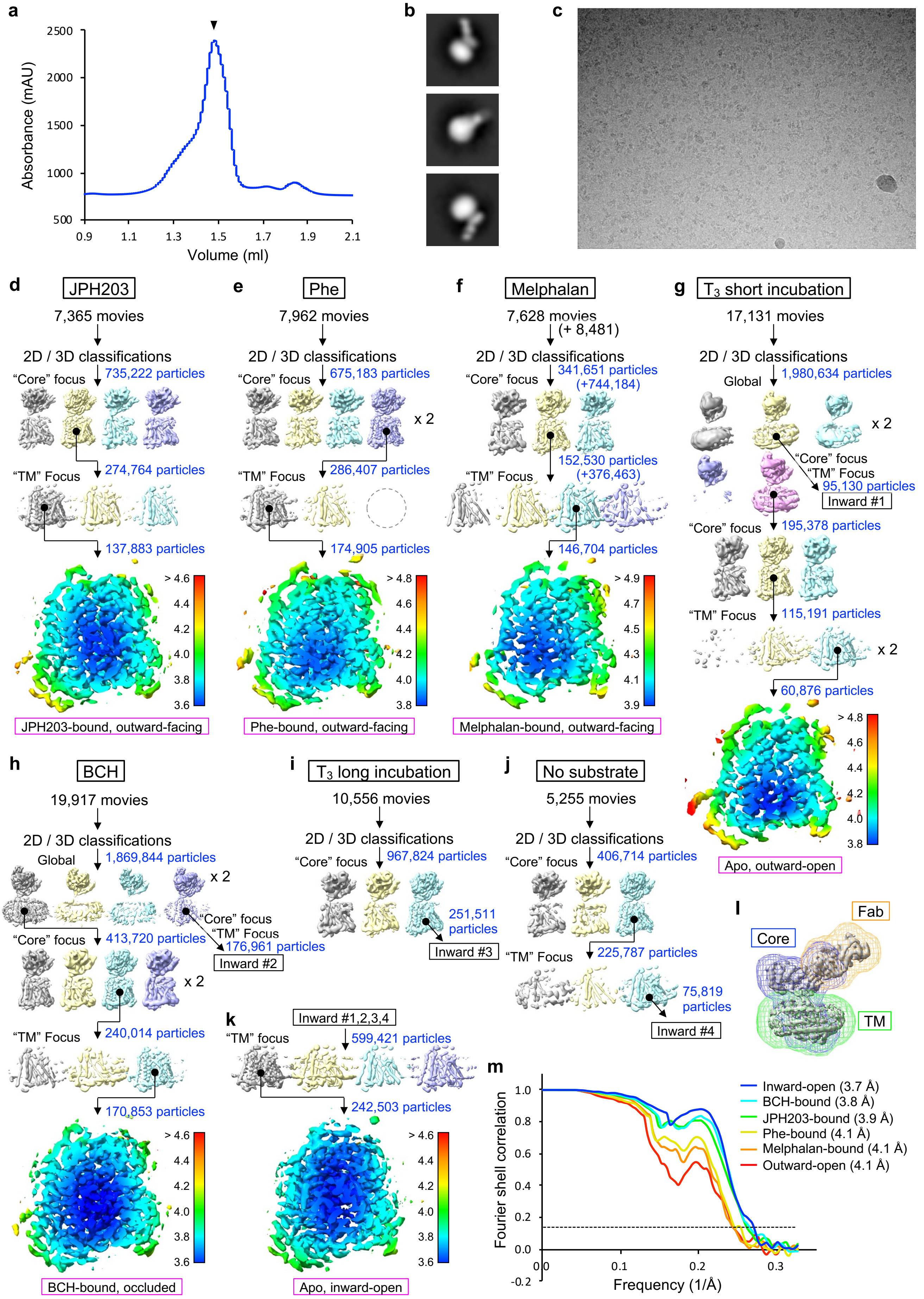


**Figure S2**

Legends on the next page

**Figure S2 | Nanodisc reconstitution, Fab screening and cryo-EM analysis of LAT1–CD98hc**
**a**) Size exclusion chromatography of LAT1–CD98hc in nanodiscs after binding Fab170.
**b**) LAT1–CD98hc + Fab170 in nanodiscs imaged by negative-stain electron microscopy. Representative 2D class averages are displayed.
**c)** Representative micrograph of LAT1–CD98hc + Fab170 bound to JPH203.
**d–j)** Single-particle processing workflows of the seven datasets used in this study.
**k)** Reconstruction of the inward-open structure by combining particles from four datasets ($T_3$ short incubation, BCH, $T_3$ long incubation and without substrate).
**l)** Masks used for multi-body refinement.
**m)** Gold-standard half-map FSC curves for the final consensus reconstruction of the six structures. The FSC was calculated based on the phase-randomization procedure as implemented in RELION.

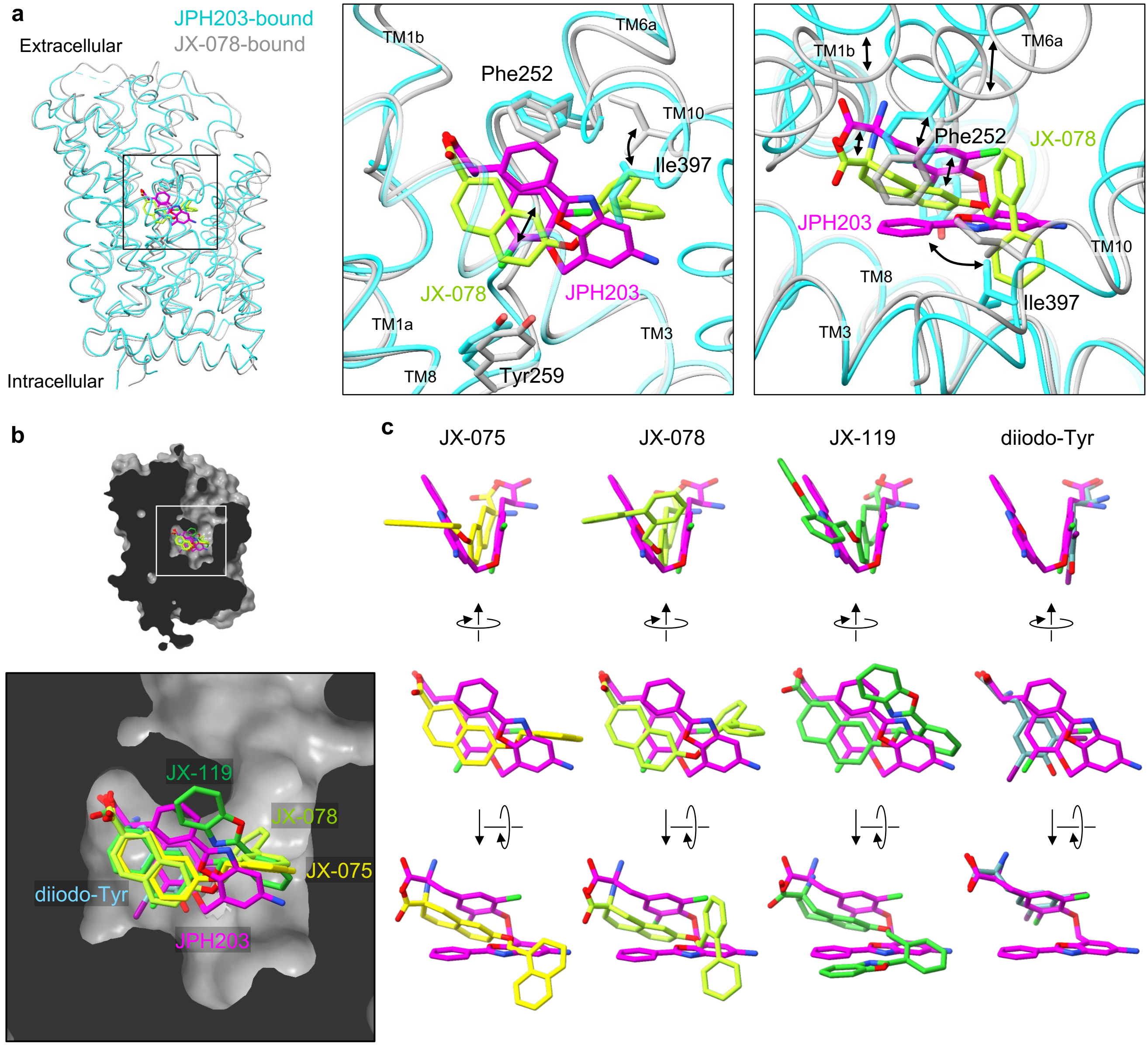


**Figure S3 | Structural comparison of JPH203 and JX inhibitors**

**a)** Overlay of two LAT1 structures bound to JPH203 or JX-078, superimposed on TM3 and TM8. Zoom-up views highlight the major structural differences of the ligand and the protein depicted as black arrows.

**b)** Superposition of JPH203, JX-075, JX-078, JX-119 and diiodo-Tyr in the substrate-binding pocket of LAT1. The structures were superposed based onTM3 and TM8. The molecular surface is shown only for the JPH203-bound structure.

**c)** Overlays of JPH203 to JX-075, JX-078, JX-119 or diiodo-Tyr observed in the pocket of LAT1.

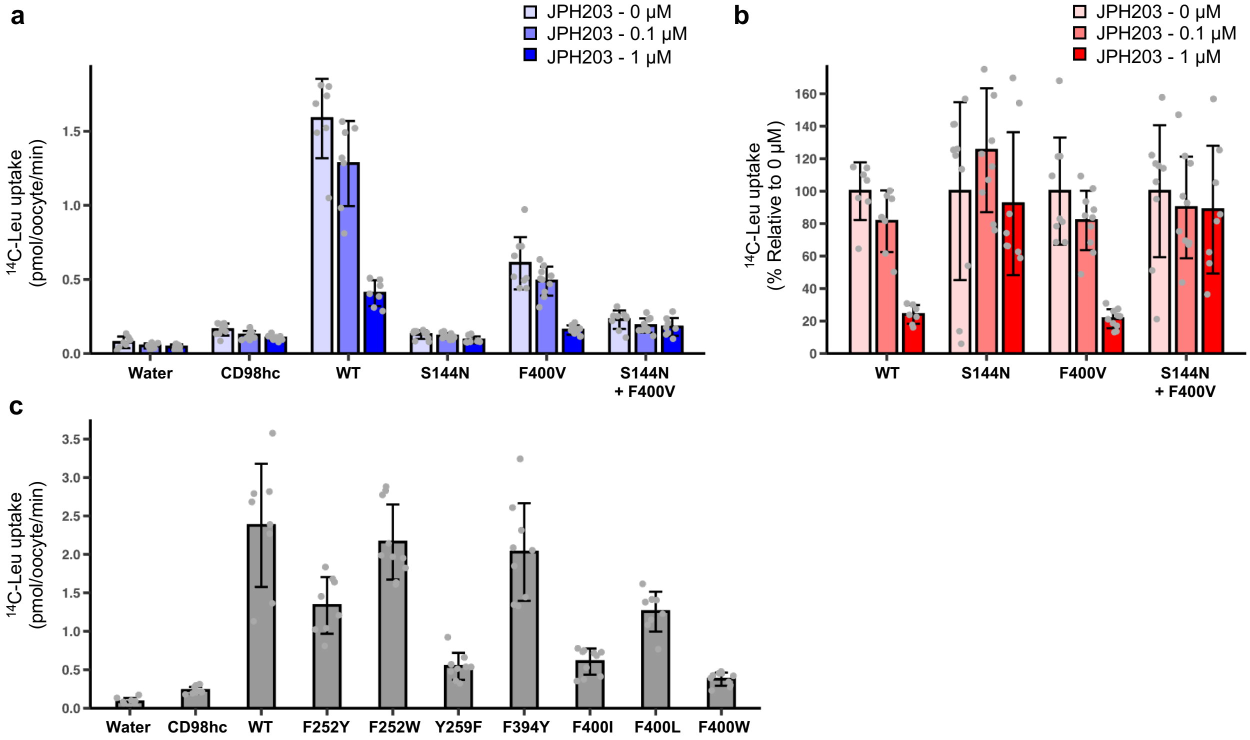


**Figure S4 | Investigating the activity and the JPH203 selectivity of LAT1 variants**

**a)** The uptake of L-[$^{14}$C]Leu into *Xenopus* oocytes expressing CD98hc and different variants of LAT1 at increasing concentrations of JPH203 in the external buffer. Data are mean ± SD and each data point represents a single oocyte (n = 7–10).

**b)** The same data as in panel **a**, plotted as % to the reference measured at 0 μM JPH203.

**c)** The activity of the LAT1 variants analyzed. Data are mean ± SD and each data point represents a single oocyte (n = 8–10).

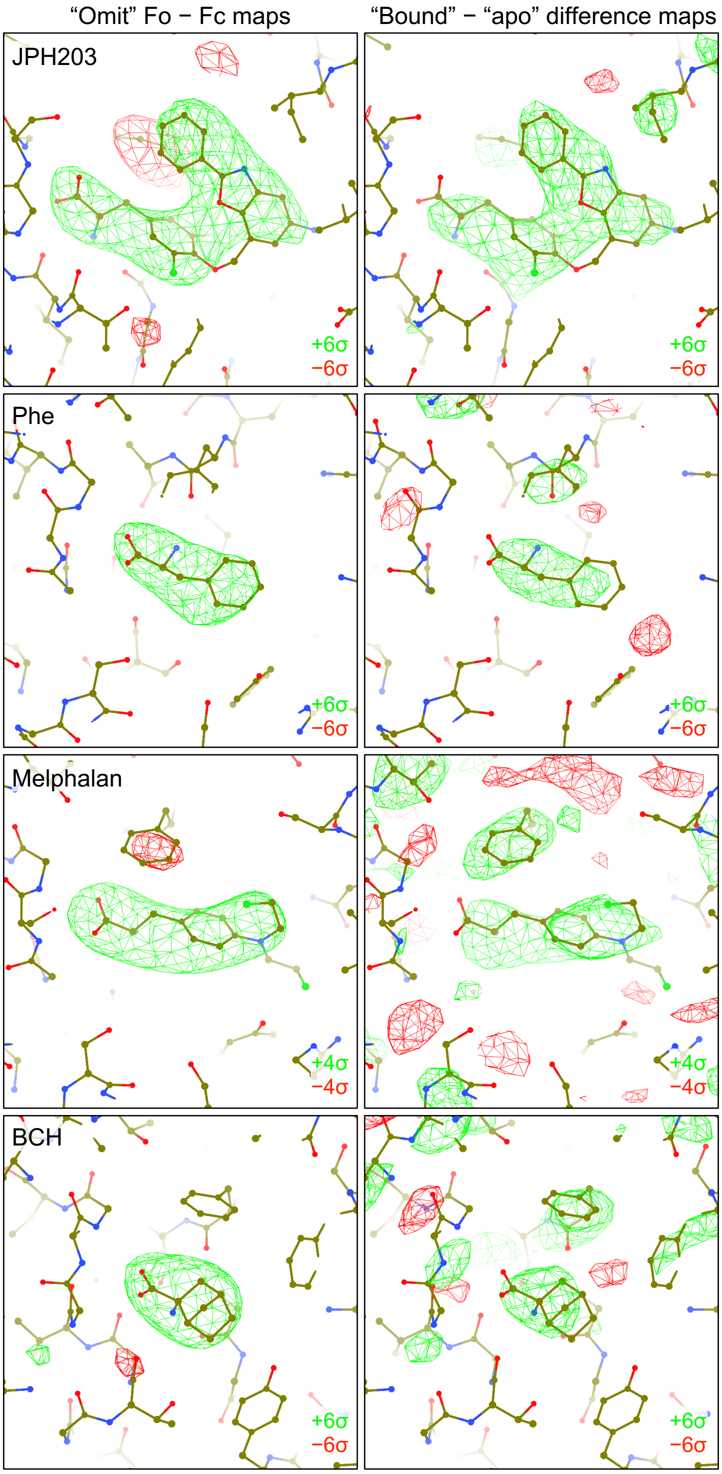


**Figure S5 | Ligand validation**

To validate the presence of ligands, difference maps were calculated using two methods. For the “omit” Fo − Fc density maps (left), the two unfiltered half-maps from RELION and the atomic model without the ligand were used as inputs for Fo − Fc difference map calculation for each ligand-bound data in servalcat. Green mesh represents positive densities and red mesh represents negative densities. For the Fo − Fo difference maps (right), the ligand-bound and apo outward-open maps were first superimposed using the likelihood-based “overlay” function in EMDA. Difference maps were then calculated for the pair of overlayed maps using the “diffmap” function. Both methods yielded strong positive densities consistent with the expected ligands, confirming the ligand-bound structures. Note that for the BCH-bound structure the difference maps reflect the structural changes between the outward-facing and occluded conformations, showing more difference peaks in the protein region.

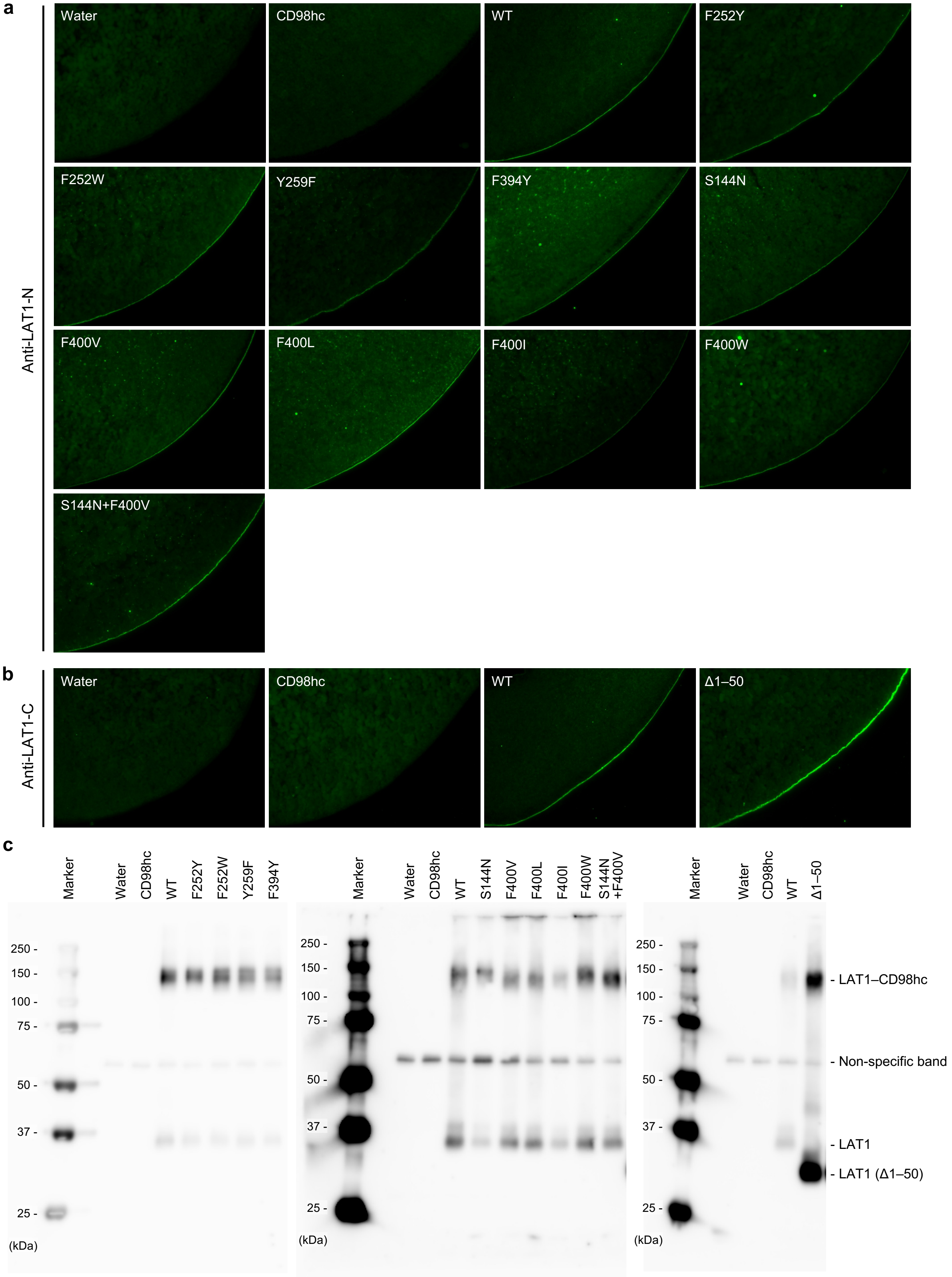


**Figure S6 | Cell-surface expression of LAT1 mutants in *Xenopus* oocytes**

**a,b)** Immunofluorescence imaging of *Xenopus* oocytes co-expressing CD98hc and different variants of LAT1, using polyclonal antibodies against the N-terminal (**a**) or C-terminal (**b**) region of LAT1.

**c)** Immunoblot analyses of membrane fractions isolated from *Xenopus* oocytes co-expressing CD98hc and different variants of LAT1, using polyclonal antibodies against the N-terminal region of LAT1.

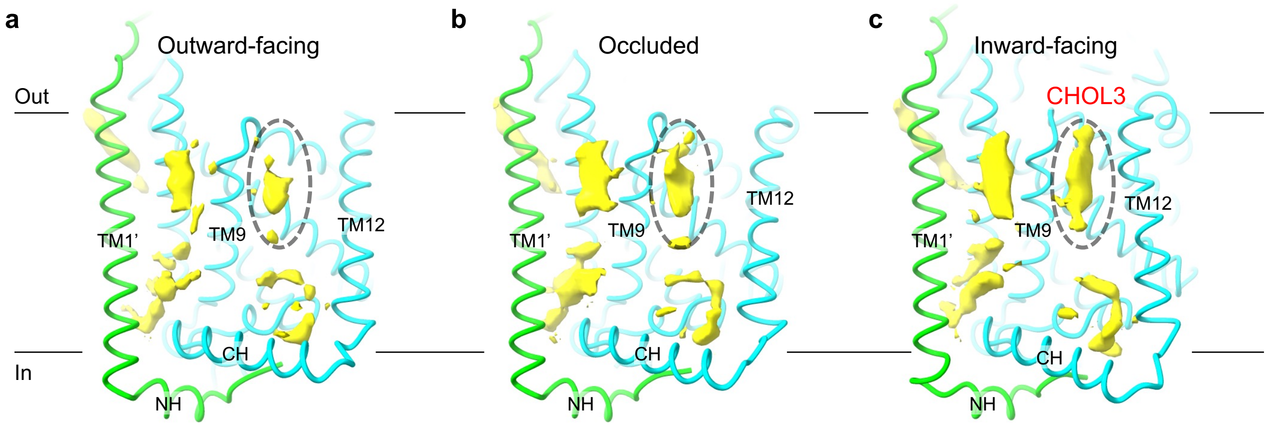


**Figure S7 | Conformation-specific lipids in LAT1**

**a–c**) Structures of the outward-facing (**a**), occluded (**b**), and inward-facing (**c**) conformations of LAT represented as tube models. All maps showed additional cryo-EM densities around the transmembrane region, most likely representing lipid molecules. One of them, located in a cleft between TM9 and TM12, showed a clear cholesterol-like shape only in the inward-facing conformation (**c**) and becomes blurred when the transporter transitions to the occluded (**b**) and outward-facing (**a**) conformations. The disappearance of the lipid density could be attributed to the dissociation of TM12 away from TM9, which reshapes the protein surface so that cholesterol can no longer bind.

### Table S1 | Cryo-EM data collection

| | JPH203 | Phe | Melphalan | $T_3$ (short) | BCH | $T_3$ (long) | No substrate |
|---|---|---|---|---|---|---|---|
| **Data collection** | | | | | | | |
| Magnification | 105,000 | 105,000 | 105,000 | 105,000 | 105,000 | 105,000 | 105,000 |
| Voltage (kV) | 300 | 300 | 300 | 300 | 300 | 300 | 300 |
| Electron exposure ($e^-/Å^2$) | 51.0 | 51.0 | 51.0 | 51.0 | 51.0 | 51.0 | 51.0 |
| Defocus range (μm) | −0.8 to −2.0 | −0.8 to −2.0 | −0.8 to −2.0 | −0.8 to −2.0 | −0.8 to −2.0 | −0.8 to −2.0 | −0.8 to −2.0 |
| Calibrated pixel size (Å) | 0.837 | 0.837 | 0.837 | 0.837 | 0.837 | 0.837 | 0.837 |
| Initial particles images (no.)[a] | 735,222 | 675,183 | 1,085,835 | 1,980,634 | 1,869,844 | 967,824 | 406,714 |

[a] After particle pre-cleaning with 2D/3D classifications

### Table S2 | Data processing, model building and validation statistics

| | JPH-bound outward-facing | | Phe-bound outward-facing | | Melphalan-bound outward-facing | | Apo outward-open | | BCH-bound occluded | | Apo inward-open | |
|---|---|---|---|---|---|---|---|---|---|---|---|---|
| **Data processing** | | | | | | | | | | | | |
| Final particle images (no.) | 137,883 | | 174,905 | | 146,704 | | 60,876 | | 170,853 | | 242,503 | |
| Final pixel size (Å) | 1.5345 | | 1.5345 | | 1.5345 | | 1.5345 | | 1.5345 | | 1.5345 | |
| Symmetry imposed | C1 | | C1 | | C1 | | C1 | | C1 | | C1 | |
| | Consensus | TMD | Consensus | TMD | Consensus | TMD | Consensus | TMD | Consensus | TMD | Consensus | TMD |
| EMDB ID | EMD-37132 | EMD-37134 | - | EMD-37140 | - | EMD-37141 | - | EMD-37142 | EMD-37135 | EMD-37136 | EMD-37137 | EMD-37138 |
| PDB ID | 8KDD | 8KDF | - | 8KDN | - | 8KDO | - | 8KDP | 8KDG | 8KDH | 8KDI | 8KDJ |
| Map resolution (Å) | | | | | | | | | | | | |
| Half map FSC = 0.143 | 3.83 | 3.89 | 3.94 | 4.12 | 3.89 | 4.12 | 3.89 | 4.12 | 3.68 | 3.78 | 3.58 | 3.73 |
| Map sharpening *B* factor ($Å^2$) | –101.5 | –138.7 | –125.6 | –196.8 | –133.2 | –184.6 | –82.2 | –145.9 | –108.4 | –163.0 | –98.4 | –171.7 |
| Local resolution range (Å) | 3.5–5.0 | 3.6–4.6 | 3.6–5.4 | 3.8–4.8 | 3.6–5.6 | 3.9–4.9 | 3.5–6.0 | 3.8–4.8 | 3.4–5.0 | 3.6–4.6 | 3.2–5.0 | 3.6–4.6 |
| **Refinement** | | | | | | | | | | | | |
| Initial model (PDB codes) | 6IRS | 6IRS | | 6IRS | | 6IRS | | 6IRS | 6IRS | 6IRS | 6IRS | 6IRS |
| Refinement resolution (Å) | 3.6 | 3.7 | | 3.9 | | 4.0 | | 4.0 | 3.6 | 3.7 | 3.6 | 3.7 |
| Map-model FSC = 0.5 (Å) | 3.6 | 3.6 | | 3.9 | | 4.0 | | 3.9 | 3.5 | 3.7 | 3.5 | 3.7 |
| Model composition | | | | | | | | | | | | |
| Non-hydrogen atoms | 10,626 | 4,045 | | 3,930 | | 3,937 | | 3,918 | 10,601 | 4,022 | 10,532 | 4,013 |
| Protein residues | 1,361 | 507 | | 506 | | 506 | | 506 | 1,361 | 506 | 1,353 | 499 |
| No. ligands | GlcNAc: 4 JPH203: 1 | CHOL: 2 POPC: 1 JPH203: 1 | | Phe: 1 | | Melphalan : 1 | | - | GlcNAc: 4 BCH: 1 | CHOL: 2 POPC: 1 BCH: 1 | GlcNAc: 4 | CHOL: 3 POPC: 2 |
| Average *B* factors ($Å^2$) | | | | | | | | | | | | |
| Protein | 211.0 | 173.9 | | 234.1 | | 226.1 | | 209.2 | 206.3 | 184.8 | 206.5 | 209.7 |
| Other | 210.1 | 192.4 | | 177.2 | | 273.4 | | - | 359.2 | 198.9 | 379.2 | 228.0 |
| R.m.s. deviation | | | | | | | | | | | | |
| Bond lengths (Å) | 0.0172 | 0.0159 | | 0.0165 | | 0.0162 | | 0.0166 | 0.177 | 0.0186 | 0.0140 | 0.0164 |
| Bond angles (°) | 2.27 | 2.14 | | 2.16 | | 2.08 | | 2.17 | 2.34 | 2.71 | 2.17 | 2.25 |
| **Validation** | | | | | | | | | | | | |
| MolProbity score | 1.75 | 1.69 | | 2.08 | | 1.88 | | 2.14 | 1.85 | 2.81 | 1.92 | 1.88 |
| Clashscore | 3.16 | 3.92 | | 6.10 | | 3.73 | | 6.24 | 2.97 | 10.57 | 3.99 | 4.09 |
| Rotamer outliers (%) | 3.27 | 2.58 | | 4.92 | | 4.23 | | 4.69 | 4.13 | 12.68 | 4.42 | 4.06 |
| Cβ outliers (%) | 0.32 | 0.00 | | 0.86 | | 0.21 | | 0.64 | 0.48 | 0.86 | 1.12 | 0.22 |
| CaBLAM outliers (%) | 1.34 | 0.60 | | 0.00 | | 0.60 | | 0.80 | 1.64 | 1.01 | 1.50 | 1.02 |
| Ramachandran plot | | | | | | | | | | | | |
| Favored (%) | 96.23 | 96.81 | | 96.60 | | 96.40 | | 95.80 | 95.78 | 93.40 | 96.35 | 96.55 |
| Allowed (%) | 3.27 | 3.19 | | 3.20 | | 3.60 | | 3.80 | 4.22 | 6.60 | 3.57 | 3.25 |
| Outliers (%) | 0.07 | 0.00 | | 0.20 | | 0.00 | | 0.40 | 0.00 | 0.00 | 0.07 | 0.20 |
